# Nucleation complex behaviour is critical for cortical microtubule array homogeneity and patterning

**DOI:** 10.1101/2022.04.05.487129

**Authors:** Bas Jacobs, René Schneider, Jaap Molenaar, Laura Filion, Eva E. Deinum

## Abstract

Plant cell walls are versatile materials that can adopt a wide range of mechanical properties through controlled deposition of cellulose fibrils. Wall integrity requires a sufficiently homogeneous fibril distribution to cope effectively with wall stresses. Additionally, specific conditions, such as the negative pressure in water transporting xylem vessels, may require more complex wall patterns, e.g., bands in protoxylem. The orientation and patterning of cellulose fibrils is guided by dynamic cortical microtubules. New microtubules are predominantly nucleated from parent microtubules causing positive feedback on local microtubule density with the potential to yield highly inhomogeneous patterns. Inhomogeneity indeed appears in all current cortical array simulations that include microtubule-based nucleation, suggesting that plant cells must possess an as-yet unknown balancing mechanism to prevent it. Here, in a combined simulation and experimental approach, we show that the naturally limited local recruitment of nucleation complexes to microtubules can counter the positive feedback, whereas local tubulin depletion cannot. We observe that nucleation complexes are preferentially inserted at microtubules. By incorporating our experimental findings in stochastic simulations, we find that the spatial behaviour of nucleation complexes delicately balances the positive feedback, such that differences in local microtubule dynamics – as in developing protoxylem – can quickly turn a homogeneous array into a patterned one. Our results provide insight into how the plant cytoskeleton is wired to meet diverse mechanical requirements and greatly increase the predictive power of computational cell biology studies.

**Significance statement:** The plant cortical microtubule array is an established model system for self-organisation, with a rich history of complementary experiments, computer simulations, and analytical theory. Understanding how array homogeneity is maintained given that new microtubules nucleate from existing microtubules has been a major hurdle for using mechanistic (simulation) models to predict future wall structures. We overcome this hurdle with detailed observations of the nucleation process from which we derive a more “natural” nucleation algorithm. With this algorithm, we enable various new lines of quantitative, mechanistic research into how cells dynamically control their cell wall properties. At a mechanistic level, moreover, this work relates to the theory on cluster coexistence in Turing-like continuum models and demonstrates its relevance for discrete stochastic entities.

## 1 Introduction

The plant cell wall is a highly versatile structure that has to adopt to diverse mechanical requirements [1, 2, 3]. Wall mechanical properties are tuned through chemical composition and, critically, through anisotropic deposition of wall material [4, 5, 6]. A key structure in this process is the cortical microtubule array, which determines where cell wall materials are inserted [7, 8, 9, 10] and guides the deposition, and hence, orientation of cellulose microfibrils, the main load-bearing component of cell walls and determinant of their anisotropic mechanical properties [7, 11, 12, 13]. The cortical array responds to various mechanical [14, 15], geometrical [16, 17, 18], developmental [19, 20, 21], and environmental [19] cues, integrating this information for future plant growth and function. This ability to respond to local wall stresses and other cues introduces a morpho-mechanical feedback loop that is considered the central ingredient of current plant growth models [22].

To make cell walls meet diverse mechanical requirements, the dynamic cortical microtubules can self-organise into various ordered structures [23], as illustrated by our focal examples: the arrays can be fully homogeneous, like the highly aligned transverse arrays of *elongating interphase cells* (Fig. 1A) [24], or locally patterned like the bands observed in developing *protoxylem* elements (Fig. 1B) [25]. Both cases require an even distribution of wall material and, therefore, of microtubules, either over the entire membrane, or among the bands. It has surfaced from multiple modelling studies [26, 27, 28, 29, 20], however, that achieving the required degree of homogeneity is far from trivial. It remains an open question how plants meet this recurring homogeneity requirement.

**Figure 1:**
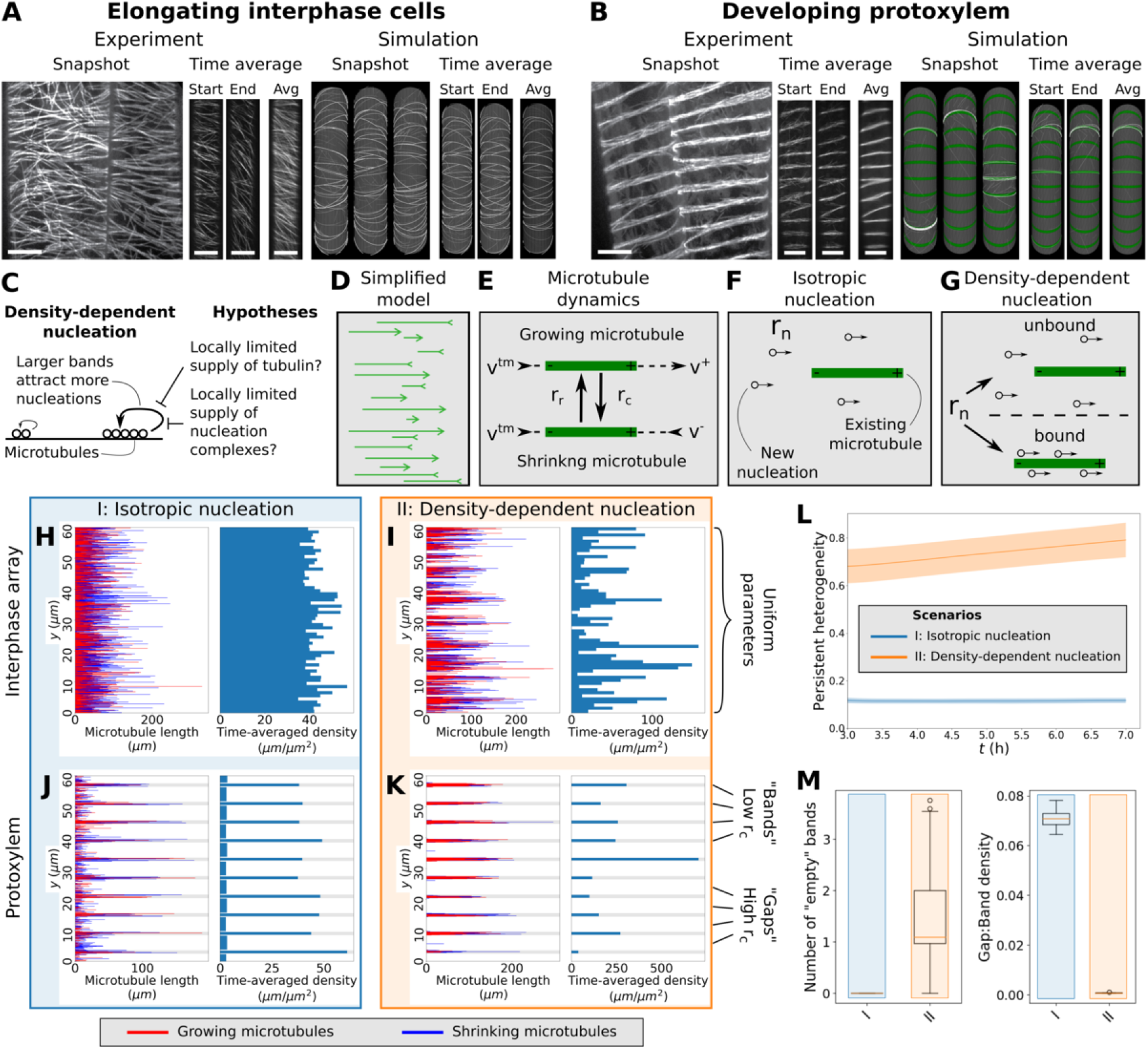
The inhomogeneity problem is reproduced by our simplified model. (A,B) Elongating interphase cells (A, left) have homogeneous arrays of transversely oriented microtubules, while simulations with density- dependent nucleation (A, right) yield highly inhomogeneous arrays. In developing protoxylem (B), microtubules are evenly distributed over a number of bands (B, left), but in simulations with density-dependent nucleation, microtubules accumulate in a small number of bands (B, right). Scale bars are 10 *µm*. (C) Hypotheses for breaking the global competition caused by density-dependent nucleation. (D–G) Implementation of the simplified model with all microtubules perfectly transversely oriented (D). Under standard microtubule dynamics (E), microtubules grow or shrink at their plus ends with rates *v*^+^ and *v*^*−*^ respectively, retract at their minus ends with rate *v*^*tm*^, and undergo spontaneous catastrophes and rescues at rates *r*_*c*_ and *r*_*r*_ respectively. Under isotropic nucleation (F), microtubules nucleate at a fixed rate *r*_*n*_ at random (*y*-)positions. With density-dependent nucleation (G), nucleations still occur at a constant global rate *r*_*n*_, but a portion of these nucleations is now distributed across existing microtubules proportional to their length. (H–K) Microtubule lengths and positions and time-averaged microtubule density of representative simulations using the simplified model for interphase arrays (H,I) and developing protoxylem (J,K). Protoxylem simulations were run for two hours without increased catastrophe rate in gaps followed by five hours with an increase of a factor three. Other simulations were run for seven hours. Time-averaging was done over the last 3 hours. (L) Measure of persistent heterogeneity over time for simulations of the interphase arrays of elongating cells. Lines and shaded areas indicate mean and standard deviation, respectively. (M) Number of empty bands (bands with less than 25% of average microtubule density in bands) and ratio of microtubule density between gaps and bands for the protoxylem simulations. Boxplots are based on quantities averaged over the last two hours of the simulations. Quantities in (L) and (M) were calculated from 100 independent simulations per nucleation mode.

The cortical microtubule array is a model system for self-organisation. A rich tradition of biophysical models [30, 31, 27, 28, 26, 32, 33, 34] heavily founded upon quantitative experiments [35, 36, 37, 38, 39] has resulted in the current consensus model for spontaneous alignment dubbed “survival-of-the-aligned” [40]. Simulation models continue to play a crucial role in understanding array behaviour, for example in the ongoing effort of unravelling how cells weigh the various and possibly conflicting cues for array orientation [19, 41, 42, 43, 44, 45, 18]. Currently, however, there are critical limitations to the application of these models as realistic simulators of the microtubule cytoskeleton, thus hampering progress on the above and other questions. The most striking shortcoming of this model is that, whenever the important aspect of microtubule bound nucleation of new microtubules is incorporated, which is experimentally observed [38], this results in highly inhomogeneous arrays (Fig. 1A,B) [26, 20, 27, 28, 29]. This, what we call, *“inhomogeneity problem”* arises because microtubule-based nucleation introduces a positive feedback that amplifies fluctuations in local microtubule density.

The existence of this positive feedback is supported by multiple experimental observations: In an established array, almost all microtubules are nucleated by *γ*-tubulin ring complexes [46, 29]. These nucleation complexes are enriched in microtubule dense regions [39], and occur almost exclusively in the microtubule bands of developing protoxylem once these are established [20]. Microtubule bound nucleation complexes, moreover, nucleate at a higher rate than unbound complexes [39]. Therefore, some mechanism must balance this positive feedback. Two likely scenarios are: 1) a local limitation of microtubule growth through the depletion of available tubulin subunits and 2) a local saturation of the amount of nucleation complexes that a microtubule-dense region can attract.

Microtubules grow through the incorporation of GTP-tubulin, mostly at their plus end. When shrinking, however, these subunits are typically released as GDP-tubulin [47, 35]. Consequently, a high local density of dynamic microtubules implies both a high consumption and release of tubulin. The two different tubulin states, however, make that tubulin is released in an inactive form, which in the very different context of small GTPase patterning provides a mechanism for the stable coexistence of multiple clusters [48]. Therefore, our first hypothesis is that local tubulin depletion could only serve as a homogenising factor when considering both tubulin states.

The second option for balancing the positive feedback and thus limiting the local increase of microtubule density may come naturally with revisiting the nucleation process. From pioneering work [31, 30, 28, 27] to current studies [33, 45, 17, 18], great progress has been made using *isotropic nucleation*, i.e., with uniform random location and orientation of new microtubules. In reality, however, most nucleations occur from nucleation complexes bound to existing microtubules, with new microtubules either parallel to their parent microtubules or branching at angles around 35° [49, 38, 39]. So far, this microtubule bound nucleation has been modelled as *density-dependent nucleation*: distributing the relevant nucleations over the existing microtubules proportional to their lengths [26, 28, 27, 29]. Density-dependent nucleation has several effects in simulations: it expands the range of biological parameters for which microtubules will spontaneously align, accelerates the alignment process in interphase arrays [26, 34], and speeds up protoxylem patterning [20]. However, this density-dependent nucleation also leads to a global competition for nucleations, in which the microtubule densest region attracts the most nucleations (Fig. 1C), resulting in a strong local clustering of microtubules in simulated interphase arrays [26, 27, 28] and many missing bands in simulated protoxylem [20] (Fig. 1B). The core of the inhomogeneity problem is that with density-dependent nucleation, nucleation sites are distributed as if the system is well-mixed, so doubling the local density somewhere will double the probability that it will attract a specific nucleation at the expense of the rest of the array. In reality, however, the docking of a nucleation complex is primarily a local process. Although increasing the local microtubule density may speed up this process, it will affect only nearby but not distant nucleation complexes, so the local increase of nucleation must saturate. By liberally extending the analogy with small GTPase patterning —where decreasing benefits of increased local density favour cluster coexistence [50]— we arrive at our second hypothesis that the locally saturating nucleation rates would suppress the global competition for nucleations and support array homogeneity.

Here, we explore the potential of our two hypotheses for solving the inhomogeneity problem using a simplified stochastic simulation model of transversely oriented dynamic microtubules. We release this model as a simulation platform called CorticalSimple. Because of their different homogeneity requirements, we use both the basic homogeneous interphase array and the banded transverse array from developing protoxylem as model systems. Both systems depend on the well-studied process of microtubule alignment into a transverse array, which we here take for granted. This simplification provides a computationally attractive environment for exploring diverse mechanisms. As nucleation complex dynamics is not sufficiently studied yet to model it properly, we perform detailed observations of nucleation complex behaviour, both under normal conditions and in sparse oryzalin-treated arrays.

This way, we discover that nucleation complexes are predominantly inserted near microtubules. Although this finding, at first glance, appears to aggravate the inhomogeneity problem, it turns out that our more natural nucleation algorithm that includes this feature allows for homogeneity, while at the same time improving the ability to form patterned arrays. Our findings pave the way for a new generation of microtubule simulation models with broad biological applications.

## 2 Results

### 2.1 A simplified model of transversely oriented microtubules reproduces the inhomogeneity problem

For solving the inhomogeneity problem, we simplified existing “full array models” (example snapshots in Fig. 1A,B) [26, 32] by taking alignment and orientation for granted. This means that all microtubules in our simulations are transversely oriented, i.e., they grow in the *x*-direction, and their positions are defined along the y-axis (Fig. 1D). The microtubules stochastically switch between growth and shrinkage (usually referred to as “catastrophe” and “rescue”), with parameters introduced in Fig. 1E and following the existing full array models. Example snapshots of the simplified arrays are shown in Fig. 1H–K. The *“interphase array”* has uniform parameters, whereas in *“protoxylem”*, the catastrophe rate *r*_*c*_ is increased in predefined gap regions after a 2-hour uniform initiation period, resulting in local destabilisation of microtubules in an existing transverse array as experimentally observed and modelled by Schneider et al. [20]. To validate our model, we use two types of microtubule nucleation from the full array models as a reference: isotropic nucleation, with uniformly distributed y-positions (Fig. 1F), and density-dependent nucleation, in which a density-dependent fraction of nucleation is “microtubule-bound”, with positions evenly distributed over all existing microtubule lengths [26] (Fig. 1G and Methods).

With isotropic nucleation, we obtained homogeneous arrays and bands of similar density, whereas with density-dependent nucleation, arrays became inhomogeneous and band density varied substantially, often leaving bands largely empty (Figures 1H–M). This validates the use of our model for studying the inhomogeneity problem.

Similar to full array simulations [20], the positive feedback inherent in density-dependent nucleation greatly enhanced the clearance of the gap regions. In the process, average microtubule density in the bands increased up to 6-fold relative to isotropic nucleation, reflecting the surface covered by band regions (Fig. 1J,K, Fig. S1).

We did observe two quantitative differences with full array protoxylem simulations. First, we observed fewer empty bands with density-dependent nucleation than in the full array simulations (Fig. 1B,K). Second, for both nucleation modes we observed much larger differences in band vs. gap density, so that the experimentally observed ∼10-fold difference between bands and gaps [20] was easily reproduced even with isotropic nucleation, matching theoretical predictions of steady state densities for non-interacting microtubules (see Supplementary Text 1.1). Two differences from the full array simulations could underlie these quantitative effects: first, microtubules cannot leave the band or gap region they were nucleated in and second, microtubule bundles –which can live longer than individual microtubules– do not occur in the simplified model.

### 2.2 Tubulin diffusion is too fast for sufficient local variation in microtubule growth velocities

To test if local tubulin depletion could solve the inhomogeneity problem, we made the microtubule growth speed dependent on the local (GTP-)tubulin concentration. Growing microtubules consumed (GTP-)tubulin, while shrinking microtubules released (GDP-)tubulin (Fig. 2A, Methods). GDP-tubulin was recharged into GTP-tubulin at a constant rate *β*.

**Figure 2:**
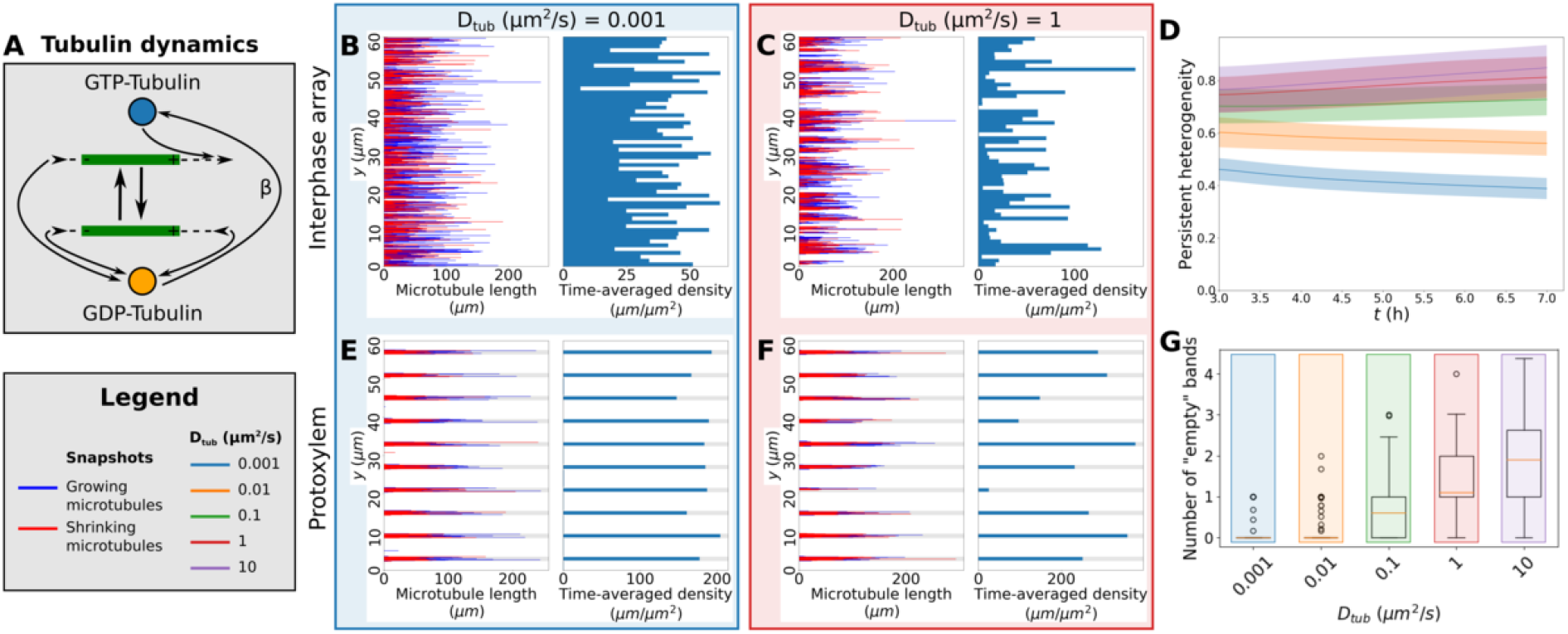
Local tubulin limitation requires unrealistically low tubulin diffusion to enhance homogeneity. (A) In our tubulin implementation, the *v*^+^ of the growing microtubules depends on the local GTP-tubulin pool, which is depleted as a result. Conversely, microtubule shrinkage increases the local GDP-tubulin concentration. GDP-tubulin is recharged at a constant rate *β* into the GTP-tubulin needed for growth. (B,C,E,F) Microtubule lengths and positions and time-averaged microtubule density of representative simulations using the simplified model with GTP- and GDP-tubulin with a tubulin recharge rate *β* = 0.01 *s*^*−*1^ and two different tubulin diffusion coefficients (*D*_*tub*_). Protoxylem simulations were run for two hours without increased catastrophe rate in gaps followed by five hours with this increase. Other simulations were run for seven hours. Time-averaging was done over the last 3 hours. (D) Measure of persistent heterogeneity over time for simulations with five different values of *D*_*tub*_. Lines and shaded areas indicate mean and standard deviation, respectively. (G) Number of empty bands (bands with less than 25% of average microtubule density in bands) for simulations with five different values of *D*_*tub*_. Boxplots are based on quantities averaged over the last two hours of the simulations. Quantities in (D) and (G) were calculated from 100 independent simulations using density-dependent nucleation.

With a plausible recharge rate of *β* = 0.01 *s*^*−*1^ [51], our simulations only produced nearly homogeneous arrays for extremely low tubulin diffusion coefficients (Fig. 2B,E), but not for values of 1 to 10 *µm*^2^*/s* similar to a measured cytoplasmic tubulin diffusion coefficient of 6 *µm*^2^*/s* [52] (Fig. 2C,D,F,G, Fig. S2).

As expected from theoretical considerations [50, 48], arrays were more homogeneous with the distinction between GTP-and GDP-tubulin than without (Fig. S3). For the lowest diffusion coefficients, tubulin essentially was a local resource over the time of the simulation, and any observed array homogeneity simply reflected the homogeneous initial tubulin distribution (Fig. S4).

These results suggest that, although the tubulin-depletion mechanism could improve homogeneity in principle, it does not ensure homogeneity in practice.

### 2.3 Nucleation complexes are preferentially inserted at microtubules

Since we found that local tubulin depletion does not solve the inhomogeneity problem, we next focused on nucleation. Little is known, however, about the mobility of nucleation complexes in the membrane itself, because normally most complexes are bound to microtubules. We, therefore, treated cells with the microtubule-depolymerising drug oryzalin [53] to reduce the density of the cortical microtubules and observed GFP-labelled *γ*-tubulin complex protein (GCP)3, a component of the nucleation complex, using spinning disc confocal microscopy. We found that microtubule-bound complexes were indeed immobile, while complexes that appeared independent of microtubules in the plasma membrane showed diffusive behaviour with a diffusion coefficient of approximately 0.013 *µm*^2^*/s* (Fig. 3A,B). Lifetimes of bound and unbound nucleation complexes were similar to those found by others [39], validating the use of these cells.

**Figure 3:**
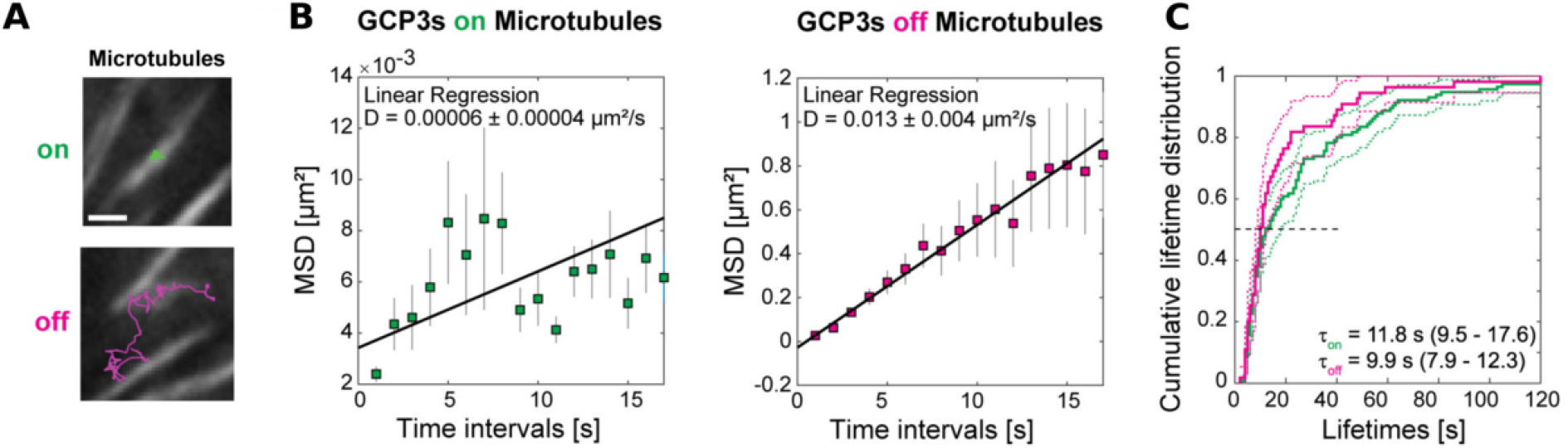
Microtubule nucleation complexes are statically bound to microtubules and diffusively anchored to plasma membranes. (A) Example trajectories of tracked GCP3 foci located on microtubules (green, top) and off microtubules (magenta, bottom). Scale bar = 1 *µm*. (B) Mean-squared displacement (MSD) calculated for 115 microtubule-bound and 55 diffusing GCP3 foci. Squares and line extensions represent means ± standard deviations for the given time intervals and the solid line represents a weighted linear fit yielding the diffusion constant and fit error. The data was recorded from nine cells and five seedlings. (C) Cumulative lifetime distributions show that the median lifetime of the tracked GCP3 foci on microtubules (green) and off microtubules (magenta) are similar. Solid and dashed lines represent the empirical cumulative distribution function and the 95% lower and upper confidence bounds for the evaluated function values, respectively. The horizontal dashed line represents the median value.

By comparing new insertions in oryzalin-treated cells to those in mock-treated controls, we discovered that nucleation complex insertion occurred preferentially near microtubules. For the control cells, we found an average insertion rate of 0.0037 *µm*^*−*2^*s*^*−*1^ (Table S1), which is similar to the 0.0045 *µm*^*−*2^*s*^*−*1^ we estimated to keep the overall nucleation rate consistent with previous simulations (see Supplementary Text 1.3). To maintain consistency, we used the latter number in our simulations. Even in the microtubule-sparse oryzalin-treated cells we found an overrepresentation of nucleation complexes on the few remaining microtubules. Nucleation complexes appeared at an average rate of 0.00026 *µm*^*−*2^*s*^*−*1^, which could be separated in a rate of 0.013 *µm*^*−*2^*s*^*−*1^ for complexes appearing near/at microtubules and 0.000085 *µm*^*−*2^*s*^*−*1^ excluding these complexes (Table S1). Because nucleation complexes co-localising with microtubules may have been inserted nearby and diffused towards them between frames, the first of these three values represents an upper bound and the last a lower bound. Still, as a conservative estimate, insertion of new complexes was at least an order of magnitude smaller in absence of microtubules. Notably, the rate of 0.013 *µm*^*−*2^*s*^*−*1^ is only 3-3.5 times higher than in a normal density array, suggesting a strong local saturation of nucleation complex insertion.

### 2.4 Local limitation of nucleation complexes can solve the inhomogeneity problem

Based on these experiments, we explicitly incorporated nucleation complexes into our simulations as particles that can associate with the membrane, move around diffusively in (effectively) two dimensions, attach to a microtubule upon encounter, and eventually either disassociate from the membrane or nucleate (Fig. 4A and Methods). With this model, we investigated the impact of differential insertion rate (*r*_*b*_ within attraction radius *R* of any microtubule, *r*_*b,min*_ otherwise) and differential nucleation rates for complexes on (*r*_*n,bound*_) or off (*r*_*n,unbound*_) microtubules [39] on array homogeneity and patterning (Fig. 4).

**Figure 4:**
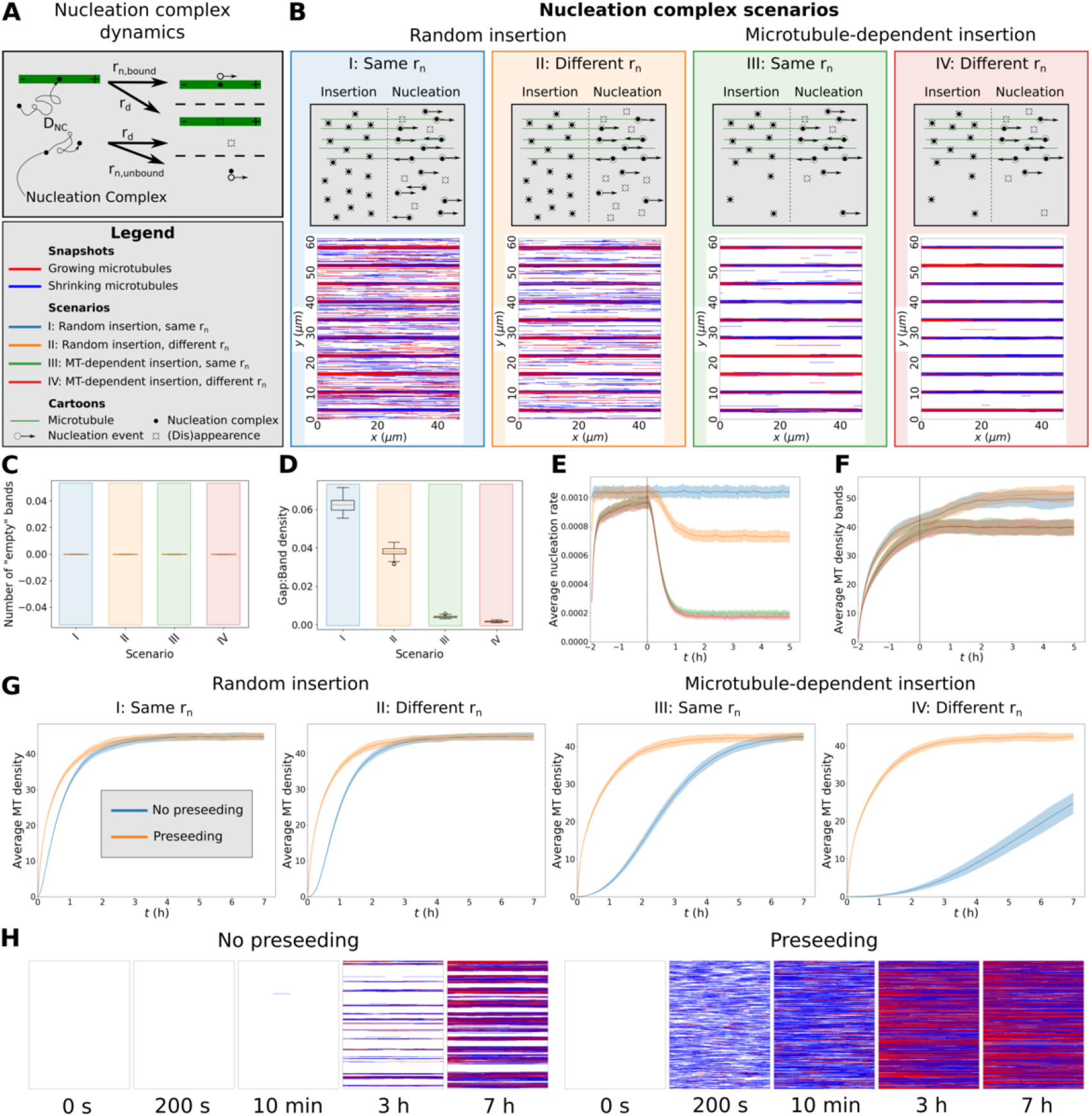
Local limitation of nucleation complex availability can ensure array homogeneity and enhance protoxylem band formation. (A) In our nucleation complex implementation, each complex moves within the two-dimensional plane of the membrane with diffusion coefficient *D*_*NC*_ and binds the first microtubule it encounters. A nucleation rate applies to each separate complex and, by default, this rate is higher for microtubule-bound complexes (*r*_*n,bound*_) than for freely diffusing (*r*_*n,unbound*_) ones. Complexes also disappear at rate *r*_*d*_. Microtubule insertion into the membrane occurs at a constant rate *r*_*b*_. For areas outside a distance *R* of a microtubule this rate is reduced to *r*_*b,min*_ in case of microtubule-dependent nucleation complex insertion. (B) Final array snapshots of representative protoxylem simulations for four different nucleation complex scenarios: I,II: Random insertion. III,IV: Microtubule-dependent insertion. I,III: No difference between bound and unbound nucleation rates. II,IV: Reduced nucleation rate for unbound complexes. (C) Number of empty bands (bands with less than 25% of average microtubule density in bands) (D) Ratio of microtubule density between gaps and bands for the protoxylem simulations. (E) Average global nucleation rates. (F) Average microtubule density in band regions. Average microtubule density for simulations without bands and gaps with and without seeded nucleations. Snapshots at various time points of a simulated array with microtubule-dependent insertion and a reduced *r*_*n*_ for unbound complexes with and without seeding. All summary statistics were calculated from 100 simulations. Lines and shaded areas indicate mean and standard deviation, respectively. Boxplots are based on quantities averaged over the last two hours of the simulations. Protoxylem simulations were run for two hours without increased catastrophe rate in gaps followed by five hours with this increase. Other simulations were run for seven hours.

We expected that our discovery of microtubule-dependent insertion, as yet another factor that favours microtubule-dense regions, would further aggravate the inhomogeneity problem.

We found, however, that with a realistic attraction radius of *R* = 50 nm, twice the width of a microtubule, fully homogeneous arrays were formed over time (Fig. 4H). This was, however, a slow process, increasing the importance of a previously reported alternative source of nucleation early during *de novo* array establishment [29], here called “seeding” (Fig. 4G,H). By halving *R*, small gaps started to appear in the array, in which nucleation complex insertion was not enhanced by microtubules, lowering the total microtubule density (Fig. S5). The diffusion coefficient *D*_*NC*_ of unbound nucleation complexes would have had to be reduced by two orders of magnitude from our experimentally observed value to achieve a similar reduction of microtubule density. With further reductions, it would become impossible to sustain a normal density array (Fig. S6).

For protoxylem patterning, we found that the differential insertion rate resulted in a more effective clearing of the gap regions than the differential nucleation rate, both for data-based rate differences (Fig. 4B,D) as well as smaller *r*_*b*_ differences (Fig. S7A) and artificially matched differences in *r*_*b*_ and *r*_*n*_ (Fig. S8). Notably, the density in band regions increased only slightly during the separation process (Fig. 4F), in stark contrast to the large increase under density-dependent nucleation (Fig. S1B,D).

Homogeneity was robustly maintained, as bands were never lost (Fig. 4B,C) except with very low diffusion coefficients (Fig. S7B). Increasing *D*_*NC*_ from low values reduced separation at most 3-fold. This observation strongly suggests that the measured value is roughly optimal for enhancing local array patterning and at the same time avoiding inhomogeneity and density loss problems (Fig. S7C,D). A similar exploration with equal insertion rate showed a window of optimal *D*_*NC*_-values for separation with a reduced (but not equal) nucleation rate for unbound complexes (Fig. S9). The default *D*_*NC*_ fell within this window for sufficiently large differences between bands and gaps only (starting from *f*_*cat*_ ≈3 − 4). This observation shows that, with sufficient (evolutionary) parameter tuning, reasonable degrees of separation could be obtained without preferential insertion at microtubules, but only with large differences in microtubule dynamics. In conclusion, differential insertion increases the patterning potential of the array and makes this regime more accessible.

## 3 Discussion

We have developed a simplified model of transversely oriented microtubules and performed quantitative experimental observations of nucleation complex dynamics to investigate how plants ensure homogeneity of their cortical microtubule array and, consequently, cell wall. With our model and experiments, we identified the natural saturation of nucleation complex recruitment to microtubule dense regions as the essential realistic mechanism for countering the positive feedbacks inherent in microtubule-based nucleation that would otherwise cause severe array inhomogeneity. The key element of this mechanism is that the competition for nucleation complexes becomes local instead of global. This effect is achieved, because the local probability of attracting a specific nucleation now saturates with local microtubule density as opposed to the linear scaling under density-dependent nucleation.

Besides the two factors already known, i.e., predominant nucleation from existing microtubules [49, 38] and reduced nucleation rate for unbound complexes [39, 29], we found that the insertion of nucleation complexes in the membrane is strongly biased towards microtubules. Together, these factors enable both completely homogeneous and complexly patterned arrays, as shown with our model systems of elongating interphase cells and developing protoxylem. Moreover, this insertion bias increases the importance of “seeding” the array with alternative (“GCP-independent”) nucleations [29] to ensure its timely establishment.

Although local tubulin depletion could not ensure homogeneity, tubulin depletion can be an important factor in limiting microtubule density at the whole cell level through changes in microtubule dynamics [40, 32]. Markedly, using parameters measured in early interphase wild type cells can result in unbound microtubule growth [54, 40], whereas with all parameter sets measured in established wild type arrays, a finite steady state microtubule density exist [37, 32, 20].

Our results demonstrate the value of our simplified model as a powerful tool for solving complex problems that can be interpreted as approximately one-dimensional. The simplification of abandoning microtubule-microtubule interactions, of course, introduces some quantitative differences with the full model, which can even increase our understanding of the real system. The largest difference is that, contrary to simulations with interacting microtubules [20], we observed strong band formation with isotropic nucleation while using the same parameters for microtubule dynamics. Two factors may underlie this difference: 1) Microtubule bundling, also without microtubule-based nucleation, increases the persistence of microtubule bundles, including those in the gap regions. Indeed, in Schneider et al. [20] it is shown that microtubule turnover is an important determinant of the band/gap-separation rate. 2) The effective nucleation rate in bands is higher in the simplified model, because all nucleations inside a band give rise to microtubules that remain inside the band. In the full array model, a substantial part of these nucleations is “lost” because the microtubules quickly grow into the gap regions. Indeed, the authors also found a much stronger degree of separation with isotropic nucleation when the nucleation rate in bands was increased. Taken together, this suggests that the aspect of co-alignment between parent and new microtubule [38] plays an important role in increasing the relevant nucleation rate inside bands and, hence, band stability. Band stability could additionally be enhanced by microtubule bundling itself.

Additional speedup of the separation process occurs if the band locations match well with density fluctuations in the (initial) microtubule array [20]. This match is likely better in reality than in current models, as gap regions are dynamically specified in the presence of the microtubule array. In metaxylem, this occurs through the small GTPase patterning protein Rho of plants 11 (AtROP11) and downstream effectors MIDD1 and Kinesin-13A [55, 56]. Various ROPs and the aforementioned effectors are also expressed during protoxylem formation [57, 58] and striated AtROP7 patterns are observed in protoxylem [59]. Results so far indicate that the corresponding microtubule patterns are not simply a readout of a ROP pattern, as changes in microtubule dynamics affect both the dynamics and outcome of the patterning process [55, 20, 60].

Notably, ROPs and other polarity factors are also indicated in the specification of the preprophase band [61], the single microtubule band that forms around the nucleus prior to cell division [62, 63], and altered microtubule dynamics are observed during its formation [37]. Together, these phenomena suggest that integrating ROP patterning and microtubule dynamics into a single simulation environment will provide mechanistic insight into many processes.

How could ROPs and microtubules sometimes produce a homogeneous pattern, like in protoand metaxylem, and sometimes a highly inhomogeneous one, like the preprophase band? Coincidentally, the literature on small GTPase patterning offers deeper insights. In the most common case, when GTPases are only interconverted between active and inactive states, the system can “phase separate” into a single cluster of active GTPases [64, 65, 50, 66]. In contrast, multiple clusters can stably coexist by (1) the addition of GTPase turnover (i.e., production and degradation) [67, 50], (2) an inactive intermediate form that cannot be reincorporated into an (active) patch immediately [48] — much like the GDP-tubulin as intermediate modelled here — or (3) an additional factor that increasingly limits the growth of active patches as they get larger, like ROP GTPase activating proteins (GAPs) that increase ROP inactivation [50]. All these options have in common that the growth of larger patches is specifically limited, and a baseline supply of raw material (inactive GTPases or nucleation of new microtubules) is guaranteed for smaller patches.

This comparison immediately stresses the significance of the uniform base insertion rate of nucleation complexes into the membrane of our model as a local supply. Our experiments on oryzalin treated cells show that nucleation complexes are indeed inserted into empty regions in the plasma membrane. We expect that rapid diffusion of cytosolic nucleation complexes, likely enhanced by cytoplasmic streaming, can ensure a relatively uniform base insertion rate. Additionally, our data suggest nucleation complexes are actively released from the membrane, because despite very different insertion rates, residence times near/at and away from microtubules are very similar (Fig. 3C), particularly for non-nucleating complexes [39]. If, however, release were governed by thermodynamic equilibrium, complexes would have remained much longer when on microtubules. Active release would contribute to the sustenance of the cytosolic pool of nucleation complexes, and hence, the base insertion rate. As multiple nucleation complex subunits contain regulatory phosphorylation sites [68], the release could be in an inactive state, which would further support homogeneity [48]. One form of density-dependent growth limitation (the conceptual equivalent of GAP proteins [50]) that is present in our model is the fact that a set of *n* isolated microtubules are more effective in capturing diffusing nucleation complexes from the membrane than a single bundle of *n* microtubules of the same length^1^. Additionally, a similar but potentially stronger effect would occur if bundling of microtubules leads to shielding of part of the binding sites for nucleation complexes, thereby specifically reducing the per length insertion rate of bundled microtubules.

The above mechanisms are not mutually exclusive and can enhance each other. Moreover, plant cells may operate close to the inhomogeneous regime of global competition as occurs with density-dependent nucleation, given the existence of inhomogeneous structures like the preprophase band. If cells are indeed close to this alternative regime, a substantial local increase in the factors that recruit nucleation complexes could over time trap a large fraction of these complexes to a specific region. The “group A” TPX2/TPXL proteins are ideal, though currently speculative, candidates for supporting preprophase band formation this way, as they are indicated in the recruitment of nucleation complexes to microtubules [69] and contain a nuclear importin domain [70], which could provide the correct perinuclear positioning of a high nucleation zone upon release. Simulations show that concentrating microtubule nucleation to the future band region can indeed reproduce a preprophase band-like structure [71].

Our novel observations of nucleation complex behaviour and the solution they provide to the inhomogeneity problem pave the way for the next generation of microtubule simulation models. Some pressing biological questions that require detailed simulations *including realistic nucleation* are: 1) How do cells integrate all the different cues affecting array orientation [14, 15, 16, 17, 18, 19, 20, 21] and resolve conflicts between them? 2) How can changes in the distribution of parent-offspring nucleation angles lead to substantial changes in cell morphology as, e.g., in the *tonneau2/fass (ton2)* mutant [72]? 3) How can a continuous interaction between ROPs and their downstream effectors on the one hand [55, 56] and the microtubule array on the other hand lead to various complex wall patterns like in proto-and metaxylem?

In summary, our work enables various new lines of quantitative, mechanistic research that will improve our understanding of how cell wall properties are dynamically controlled.

## Supporting information

all supplementary material

## 4 Acknowledgements

R.S. would like to acknowledge Deutsche Forschungsgemeinschaft grant 453188536.

## 5 Methods

### 5.1 Simulation setup

Cortical microtubules exist effectively on a two-dimensional surface on the inside of the membrane. Therefore, we chose our simulation domain to represent the cortex of a cylindrical cell, with a height of 60 *µm* and a radius of 7.5 *µm* as in Schneider et al. [20]. This domain was represented by a rectangle with a horizontal periodic *x*-axis and a vertical *y*-axis representing the circumference and length of the cylinder, respectively. Except for the variant with nucleation complexes, horizontal positions were irrelevant and not tracked in the simulations. CorticalSimple is written in python and can be downloaded from git.wur.nl/Biometris/ articles/corticalsimple

### 5.2 Core microtubule dynamics

Microtubule growth, shrinkage, and minus end retraction (treadmilling) occur at speeds *v*^+^, *v*^*−*^, and *v*^*tm*^, respectively, with catastrophes and rescues occurring at rates *r*_*c*_ and *r*_*r*_, respectively (Fig. 1E). These are the same dynamics as described by Tindemans et al. [32] and parameters were based on [20] (see Table S2 for parameter values).

### 5.3 Protoxylem band and gap regions

Protoxylem simulations used ten band regions of 1 *µm* separated by gap regions of 5 *µm* (with 2.5 *µm* gap regions on either end of the domain). Following [20], microtubule stability was reduced in the gap regions by increasing the catastrophe rate by a factor *f*_*cat*_ (default: *f*_*cat*_ = 3) compared to the band regions. Before this increase in gap catastrophe rate, simulations were run for two hours with homogeneous parameters (*f*_*cat*_ = 1), allowing a microtubule array to form.

### 5.4 Basic nucleation modes

Isotropic nucleations were drawn at a constant global rate of *r*_*n*_ *· A*, where *r*_*n*_ is the nucleation rate in *µm*^*−*2^*s*^*−*1^ and *A* is the domain area in *µm*^2^ and given uniformly distributed *y*-positions (Fig. 1F).

Density-dependent nucleation was implemented as in Deinum et al. [26]. Nucleation events were scheduled with a total rate *r*_*n*_, of which a density-dependent fraction was assigned to microtubules, resulting in a bound rate *r*_*n,bound*_ following:

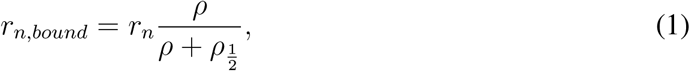

where *ρ* is the global microtubule density in *µm* microtubule per *µm*^2^ and 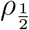 is the micro-tubule density at which half of all nucleations are bound. We then assigned *y*-positions to the unbound nucleations, as described for isotropic nucleation. The bound nucleations were distributed randomly across the total microtubule length and then got the *y*-position of their parent microtubule with a small normally distributed displacement (*σ* = 0.1*µm*), which was redrawn for positions falling outside the simulation domain (Fig. 1G).

### 5.5 Tubulin dynamics

In one dimension, tubulin dynamics for a single tubulin follows the diffusion equation:

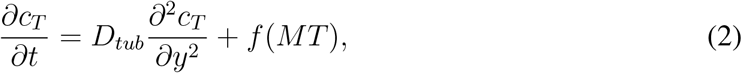

where *c*_*T*_ is the tubulin concentration, *t* is the time, *y* is the position along the longitudinal axis, *D*_*tub*_ is the tubulin diffusion coefficient, and *f* (*MT*) the function of microtubule dynamics that specifies the net release of tubulin from microtubules. This last term can be calculated directly from local changes in microtubule lengths at each integration time step. Similarly, when distinguishing GTP-and GDP-tubulin, we have:

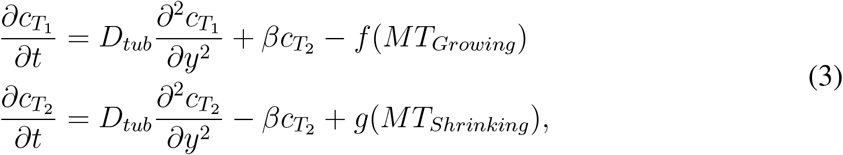

where 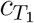 and 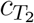 are the concentrations of GTP-and GDP-tubulin, respectively, *β* is the recharge rate at which GDP-tubulin is converted back into GTP-tubulin, and *f* (*MT*_*Growing*_) and *g*(*MT*_*Shrinking*_) are the functions of microtubule dynamics that determine the tubulin consumption by growing microtubules and the tubulin release by shrinking microtubules, respectively. For convenience, we express tubulin concentrations in *µm* of microtubule length equivalent per *µm*^2^.

Arrays were initiated without microtubules and with a uniform (GTP-)tubulin concentration of *L*_*max*_*/A*, where *A* is the domain area and *L*_*max*_ is the maximum total microtubule length when all tubulin is in microtubule form. Growth speed *v*^+^ was made linearly dependent on the local (GTP-)tubulin concentration, according to:

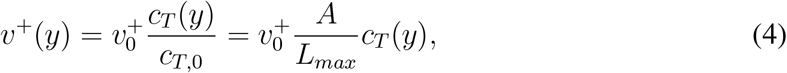

where 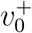 is the initial growth speed and parameter *c*_*T*,0_ is the initial homogeneous (GTP-)tubulin concentration.

The diffusion equations were integrated using a Crank-Nicolson algorithm [73] with integration steps of 0.01 *s* in time and 0.2 *µm* in space. Microtubule growth speeds were adjusted to the new tubulin concentration every time step and kept constant inbetween.

### 5.6 Nucleation complexes

In our nucleation complex implementation, membrane-associated complexes diffuse with diffusion coefficient *D*_*NC*_. If a complex runs into a microtubule, it binds the microtubule and remains stationary. Therefore, to allow complexes to pass around the ends of microtubules, the *x*-positions and microtubule directions are tracked for this model variant.

Nucleation complexes are inserted at a constant rate *r*_*b*_, which can be reduced to *r*_*b,min*_ for regions without a microtubule within a distance *R* in case of microtubule-dependent nucleation complex insertion.

Each individual complex can disassociate from the membrane at a rate *r*_*d*_ and nucleate at a rate *r*_*n*_ (Fig. 4A). Based on experimental data from Nakamura et al. [39] this nucleation rate is set a factor fifteen larger for microtubule-bound complexes than for unbound complexes (Supplementary Text 1.3). Upon nucleation, the complex involved is removed from the simulation, since experimental observations indicate that complexes that nucleated hardly ever nucleate a second time before disappearing [39]. Positions of new microtubules are adopted from the parent nucleation complex, with the same small vertical displacement that is also used for density-dependent nucleation. The growing plus-end of each new microtubule is oriented either to the left or to the right with an equal probability.

Seeded nucleations are implemented by starting simulations with nucleation complexes with uniformly distributed complex-independent nucleations at a density of 1 nucleation per *µm*^2^. This value has been chosen to be close to the steady state microtubule density.

Nucleation complex diffusion has been implemented using two-dimensional Brownian motion simulations. Complexes move independently in both horizontal and vertical directions every time step by a distance of 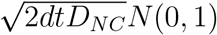, where *dt* is the time step (0.01 *s* in our simulations), and *N* (0, 1) is a standard normally distributed random number.

### 5.7 Model parameters

All simulation parameters are given in Table S2. Basic model parameters were chosen to be consistent with previous simulations of microtubule dynamics in protoxylem development [20]. In the tubulin simulations, *D*_*tub*_ and *β* were varied to study their effect. *L*_*max*_ and 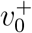 were tuned such that the average microtubule growth speed at steady state would be approximately equal to that used in simulations without tubulin (see Supplementary Text 1.2). Parameters *r*_*b*_, *r*_*d*_, *r*_*n,bound*_, and *r*_*n,unbound*_ were estimated from data by Nakamura et al. [39] (see Supplementary Text 1.3). For simulations with microtubule-dependent nucleation complex insertion, we tried several values of *r*_*b,min*_ based on our experimental measurements and chose a distance *R* of 0.05 *µm*, which is about twice the width of a microtubule.

### 5.8 Measures of heterogeneity

For protoxylem simulations, we counted the number of largely empty bands, defined as bands with less than 25% of the average microtubule density in bands. The persistent heterogeneity measure for transverse interphase arrays was calculated as the standard deviation of the values from a time-averaged microtubule density histogram divided by the average density. We used a histogram bin size of 1 *µm* and a time average over the last three hours, with one measurement every 200 s.

### 5.9 Experimental measurements

#### 5.9.1 Plant material and growth conditions

Arabidopsis (*Arabidopsis thaliana*, Columbia Col-0 ecotype) expressing the 35S promoter-driven VND7-VP16-GR (VND7) construct, the 35S promoter-driven mCHERRY-TUA5 microtubule marker as well as the GCP3-GFP microtubule nucleation marker were used [20]. Seeds were surface-sterilised and grown similar to [20]. To depolymerize MTs in epidermal cells, we applied 40*µ*M oryzalin for 4 hours and subsequently focussed on cells that showed remaining MT polymers in their cortex.

#### 5.9.2 Induction of protoxylem formation

Three-day old dark-grown seedlings were transferred to half-strength MS, 1% sucrose plates supplemented with 10*µM* dexamethasone (DEX). The unwrapped plates were then kept in the same phytotron for 24 hours. Subsequently, seedlings were transferred to a microscope slide for imaging.

#### 5.9.3 Spinning disk microscopy

Imaging was performed similar to [20]. Briefly, we used a spinning disc microscope consisting of a CSU-X1 spinning disk head (Yokogawa), an Eclipse TI (Nikon) inverted microscope body equipped with a perfect focus system, an Evolve CCD camera (Photometrics), and a CFI Apo TIRF 100x oil-immersion objective.

#### 5.9.4 Image analysis

Confocal z-stack recordings of microtubules in non-induced and VND7-induced conditions were acquired (0.3*µm* z-steps, 6*µm* z-depth, 300*ms* integration time) and surface-projected using a custom-made Matlab code. Confocal single-plane time-lapse recordings of microtubules and GCP3 foci were acquired (1 second intervals, 2 minute duration, 300ms and 500ms integration for microtubules and GCP3, respectively) and analysed using the open-source tracking software FIESTA [74]. The in-built mean-squared displacement function was used to measure the diffusion constant of GCP3 foci. Cumulative lifetime distributions were made using the in-built Matlab function ecdf.m.

## Footnotes

1 E.g., assuming a membrane residence time of 10 s, a nucleation complex would have an average diffusion length of 361 nm. For aligned same length microtubules, this reduces to a 1D problem, with a 2*361 + 10*25 nm cross section covered by a bundle of 10 microtubules and a 10*(2*361 + 25) nm cross section for the isolated microtubules. So, as a lower bound, the bundle would be only 13% as effective in capturing free nucleation complexes.

## Notes

### Competing Interest Statement

The authors have declared no competing interest.

## References

[1] Tobias I. Baskin. Anisotropic expansion of the plant cell wall. Annual Review of Cell and Developmental Biology, 21(1):203–222, 2005. PMID: 16212493.

[2] Arezki Boudaoud. An introduction to the mechanics of morphogenesis for plant biologists. Trends in plant science, 15(6):353–360, 2010.

[3] Aleksandra Sapala, Adam Runions, Anne-Lise Routier-Kierzkowska, Mainak Das Gupta, Lilan Hong, Hugo Hofhuis, Stéphane Verger, Gabriella Mosca, Chun-Biu Li, Angela Hay, et al. Why plants make puzzle cells, and how their shape emerges. Elife, 7:e32794, 2018.

[4] Siobhan A Braybrook and Henrik Jönsson. Shifting foundations: the mechanical cell wall and development. Current opinion in plant biology, 29:115–120, 2016.

[5] Gabriel Levesque-Tremblay, Jerome Pelloux, Siobhan A Braybrook, and Kerstin Müller. Tuning of pectin methylesterification: consequences for cell wall biomechanics and development. Planta, 242(4):791–811, 2015.

[6] Michael Ogden, Rainer Hoefgen, Ute Roessner, Staffan Persson, and Ghazanfar Abbas Khan. Feeding the walls: how does nutrient availability regulate cell wall composition? International journal of molecular sciences, 19(9):2691, 2018.

[7] Ryan Gutierrez, Jelmer J. Lindeboom, Alex R. Paredez, Anne Mie C. Emons, and David W. Ehrhardt. Arabidopsis cortical microtubules position cellulose synthase delivery to the plasma membrane and interact with cellulose synthase trafficking compartments. Nature Cell Biology, 11(7):797–806, July 2009.

[8] Y. Watanabe, M. J. Meents, L. M. McDonnell, S. Barkwill, A. Sampathkumar, H. N. Cartwright, T. Demura, D. W. Ehrhardt, A.L. Samuels, and S. D. Mansfield. Visualization of cellulose synthases in Arabidopsis secondary cell walls. Science, 350(6257):198–203, 2015.

[9] Rene Schneider, Lu Tang, Edwin R. Lampugnani, Sarah Barkwill, Rahul Lathe, Yi Zhang, Heather E. McFarlane, Edouard Pesquet, Totte Niittyla, Shawn D. Mansfield, Yihua Zhou, and Staffan Persson. Two complementary mechanisms underpin cell wall patterning during xylem vessel development. The Plant Cell, 29(10):2433–2449, 2017.

[10] Nemanja Vukašinović, Yoshihisa Oda, Přemysl Pejchar, Lukáš Synek, Tamara Pečenková, Anamika Rawat, Juraj Sekereš, Martin Potocký, and Viktor Žárský. Microtubule-dependent targeting of the exocyst complex is necessary for xylem development in Arabidopsis. New Phytologist, 213(3):1052–1067, 2017. 2016-22430.

[11] Alexander R. Paredez, Christopher R. Somerville, and David W. Ehrhardt. Visualization of cellulose synthase demonstrates functional association with microtubules. Science, 312(5779):1491–1495, 2006.

[12] Jordi Chan and Enrico Coen. Interaction between autonomous and microtubule guidance systems controls cellulose synthase trajectories. Current Biology, 30(5):941–947, 2020.

[13] René Schneider, Tobias Hanak, Staffan Persson, and Christian A Voigt. Cellulose and callose synthesis and organization in focus, what’s new? Current Opinion in Plant Biology, 34:9–16, 2016.

[14] Olivier Hamant, Marcus G. Heisler, Henrik Jönsson, Pawel Krupinski, Magalie Uyttewaal, Plamen Bokov, Francis Corson, Patrik Sahlin, Arezki Boudaoud, Elliot M. Meyerowitz, Yves Couder, and Jan Traas. Developmental patterning by mechanical signals in arabidopsis. Science, 322(5908):1650–1655, 2008.

[15] Leia Colin, Antoine Chevallier, Satoru Tsugawa, Florian Gacon, Christophe Godin, Virgile Viasnoff, Timothy E. Saunders, and Olivier Hamant. Cortical tension overrides geometrical cues to orient microtubules in confined protoplasts. Proceedings of the National Academy of Sciences, 117(51):32731–32738, 2020.

[16] Chris Ambrose, Jun F. Allard, Eric N. Cytrynbaum, and Geoffrey O. Wasteneys. A CLASP-modulated cell edge barrier mechanism drives cell-wide cortical microtubule organization in Arabidopsis. Nature Communications, 2:430, August 2011.

[17] Bandan Chakrabortty, Ikram Blilou, Ben Scheres, and Bela M. Mulder. A computational framework for cortical microtubule dynamics in realistically shaped plant cells. PLOS Computational Biology, 14(2):1–26, 02 2018.

[18] Pauline Durand-Smet, Tamsin A. Spelman, Elliot M. Meyerowitz, and Henrik Jönsson. Cytoskeletal organization in isolated plant cells under geometry control. Proceedings of the National Academy of Sciences, 117(29):17399–17408, 2020.

[19] Jelmer J. Lindeboom, Masayoshi Nakamura, Anneke Hibbel, Kostya Shundyak, Ryan Gutierrez, Tijs Ketelaar, Anne Mie C. Emons, Bela M. Mulder, Viktor Kirik, and David W. Ehrhardt. A mechanism for reorientation of cortical microtubule arrays driven by microtubule severing. Science, 342(6163):1245533, 2013.

[20] René Schneider, Kris van’t Klooster, Kelsey L. Picard, Jasper van der Gucht, Taku Demura, Marcel Janson, Arun Sampathkumar, Eva E. Deinum, Tijs Ketelaar, and Staffan Persson. Long-term single-cell imaging and simulations of microtubules reveal principles behind wall patterning during proto-xylem development. Nature Communications, 12(1):669, 2021.

[21] Laura Vineyard, Andrew Elliott, Sonia Dhingra, Jessica R. Lucas, and Sidney L. Shaw. Progressive transverse microtubule array organization in hormone-induced arabidopsis hypocotyl cells. Plant Cell, 25(2):662–676, Feb 2013.

[22] Duy-Chi Trinh, Juan Alonso-Serra, Mariko Asaoka, Leia Colin, Matthieu Cortes, Alice Malivert, Shogo Takatani, Feng Zhao, Jan Traas, Christophe Trehin, et al. How mechanical forces shape plant organs. Current Biology, 31(3):R143–R159, 2021.

[23] Peter Nick. Plant microtubules: development and flexibility, volume 11. Springer Science & Business Media, 2008.

[24] David W. Ehrhardt and Sidney L. Shaw. Microtubule dynamics and organization in the plant cortical array. Annual Review of Plant Biology, 57(1):859–875, 2006. PMID: 16669785.

[25] Yoshihisa Oda and Hiroo Fukuda. Secondary cell wall patterning during xylem differentiation. Current Opinion in Plant Biology, 15(1):38 – 44, 2012. Growth and development.

[26] Eva E Deinum, Simon H Tindemans, and Bela M Mulder. Taking directions: the role of microtubule-bound nucleation in the self-organization of the plant cortical array. Physical Biology, 8(5):056002, 2011.

[27] Jun F. Allard, Geoffrey O. Wasteneys, and Eric N. Cytrynbaum. Mechanisms of self-organization of cortical microtubules in plants revealed by computational simulations. Molecular Biology of the Cell, 21(2):278–286, 2010.

[28] Ezgi Can Eren, Ram Dixit, and Natarajan Gautam. A three-dimensional computer simulation model reveals the mechanisms for self-organization of plant cortical microtubules into oblique arrays. Molecular Biology of the Cell, 21(15):2674–2684, 2010. PMID: 20519434.

[29] Jelmer J. Lindeboom, Antonios Lioutas, Eva E. Deinum, Simon H. Tindemans, David W. Ehrhardt, Anne Mie C. Emons, Jan W. Vos, and Bela M. Mulder. Cortical microtubule arrays are initiated from a nonrandom prepattern driven by atypical microtubule initiation. Plant Physiology, 161(3):1189–1201, 2013.

[30] Simon H. Tindemans, Rhoda J. Hawkins, and Bela M. Mulder. Survival of the aligned: Ordering of the plant cortical microtubule array. Physical Review Letters, 104:058103, Feb 2010.

[31] Rhoda J. Hawkins, Simon H. Tindemans, and Bela M. Mulder. Model for the orientational ordering of the plant microtubule cortical array. Physical Review E, 82:011911, Jul 2010.

[32] Simon Tindemans, Eva Deinum, Jelmer Lindeboom, and Bela Mulder. Efficient event-driven simulations shed new light on microtubule organization in the plant cortical array. Frontiers in Physics, 2:19, 2014.

[33] Eva E Deinum, Simon H Tindemans, Jelmer J Lindeboom, and Bela M Mulder. How selective severing by katanin promotes order in the plant cortical microtubule array. Proceedings of the National Academy of Sciences, 114(27):6942–6947, 2017.

[34] Panayiotis Foteinopoulos and Bela M Mulder. The effect of anisotropic microtubule-bound nucleations on ordering in the plant cortical array. Bulletin of mathematical biology, 76(11):2907–2922, 2014.

[35] Sidney L. Shaw, Roheena Kamyar, and David W. Ehrhardt. Sustained microtubule tread-milling in Arabidopsis cortical arrays. Science, 300(5626):1715–1718, 2003.

[36] Ram Dixit and Richard Cyr. Encounters between dynamic cortical microtubules promote ordering of the cortical array through angle-dependent modifications of microtubule behavior. The Plant Cell Online, 16(12):3274–3284, 2004.

[37] Jan W. Vos, Marileen Dogterom, and Anne Mie C. Emons. Microtubules become more dynamic but not shorter during preprophase band formation: A possible “search-and-capture” mechanism for microtubule translocation. Cell Motility and the Cytoskeleton, 57(4):246–258, 2004.

[38] Jordi Chan, Adrian Sambade, Grant Calder, and Clive Lloyd. Arabidopsis cortical microtubules are initiated along, as well as branching from, existing microtubules. The Plant Cell Online, 21(8):2298–2306, 2009.

[39] Masayoshi Nakamura, David W. Ehrhardt, and Takashi Hashimoto. Microtubule and katanin-dependent dynamics of microtubule nucleation complexes in the acentrosomal Arabidopsis cortical array. Nature Cell Biology, 12(11):1064–1070, November 2010.

[40] Eva E Deinum and Bela M Mulder. Modelling the role of microtubules in plant cell morphology. Current Opinion in Plant Biology, 16(6):688 – 692, 2013. Cell biology.

[41] Bandan Chakrabortty, Viola Willemsen, Thijs de Zeeuw, Che-Yang Liao, Dolf Weijers, Bela Mulder, and Ben Scheres. A plausible microtubule-based mechanism for cell division orientation in plant embryogenesis. Current Biology, 28(19):3031–3043.e2, October 2018.

[42] Masayoshi Nakamura, Jelmer J Lindeboom, Marco Saltini, Bela M Mulder, and David W Ehrhardt. Spr2 protects minus ends to promote severing and reorientation of plant cortical microtubule arrays. J Cell Biol, 217(3):915–927, 2018.

[43] Marco Saltini and Bela M. Mulder. A plausible mechanism for longitudinal lock-in of the plant cortical microtubule array after light-induced reorientation. Quantitative Plant Biology, 2:e9, 2021.

[44] Marco Saltini and Bela M Mulder. Critical threshold for microtubule amplification through templated severing. Physical Review E, 101(5):052405, 2020.

[45] Vincent Mirabet, Pawel Krupinski, Olivier Hamant, Elliot M. Meyerowitz, Henrik Jönsson, and Arezki Boudaoud. The self-organization of plant microtubules inside the cell volume yields their cortical localization, stable alignment, and sensitivity to external cues. PLOS Computational Biology, 14(2):1–23, 02 2018.

[46] Takashi Hashimoto. A ring for all: γ-tubulin-containing nucleation complexes in acentrosomal plant microtubule arrays. Current Opinion in Plant Biology, 16(6):698–703, 2013. Cell biology.

[47] Arshad Desai,, and Timothy J. Mitchison. Microtubule polymerization dynamics. Annual Review of Cell and Developmental Biology, 13(1):83–117, 1997. PMID: 9442869.

[48] Jian-geng Chiou, Kyle D Moran, and Daniel J Lew. How cells determine the number of polarity sites. eLife, 10:e58768, apr 2021.

[49] Takashi Murata, Seiji Sonobe, Tobias I. Baskin, Susumu Hyodo, Seiichiro Hasezawa, Toshiyuki Nagata, Tetsuya Horio, and Mitsuyasu Hasebe. Microtubule-dependent microtubule nucleation based on recruitment of γ-tubulin in higher plants. Nature Cell Biology, 7(10):961–968, October 2005.

[50] Bas Jacobs, Jaap Molenaar, and Eva E. Deinum. Small GTPase patterning: How to stabilise cluster coexistence. PLOS ONE, 14(3):1–28, 03 2019.

[51] B J Terry and D L Purich. Nucleotide release from tubulin and nucleoside-5’-diphosphate kinase action in microtubule assembly. Journal of Biological Chemistry, 254(19):9469– 9476, 1979.

[52] E D Salmon, W M Saxton, R J Leslie, M L Karow, and J R McIntosh. Diffusion coefficient of fluorescein-labeled tubulin in the cytoplasm of embryonic cells of a sea urchin: video image analysis of fluorescence redistribution after photobleaching. Journal of Cell Biology, 99(6):2157–2164, 12 1984.

[53] LC Morejohn, TE Bureau, J Mole-Bajer, AS Bajer, and DE Fosket. Oryzalin, a dinitroaniline herbicide, binds to plant tubulin and inhibits microtubule polymerization in vitro. Planta, 172(2):252–264, 1987.

[54] Eiko Kawamura and Geoffrey O Wasteneys. Mor1, the arabidopsis thaliana homologue of xenopus map215, promotes rapid growth and shrinkage, and suppresses the pausing of microtubules in vivo. Journal of cell science, 121(24):4114–4123, 2008.

[55] Yoshihisa Oda and Hiroo Fukuda. Initiation of cell wall pattern by a rho- and microtubule-driven symmetry breaking. Science, 337(6100):1333–1336, 2012.

[56] Yoshihisa Oda and Hiroo Fukuda. Rho of plant GTPase signaling regulates the behavior of Arabidopsis kinesin-13A to establish secondary cell wall patterns. The Plant Cell Online, 25(11):4439–4450, 2013.

[57] Masatoshi Yamaguchi, Nobutaka Mitsuda, Misato Ohtani, Masaru Ohme-Takagi, Ko Kato, and Taku Demura. VASCULAR-RELATED NAC-DOMAIN 7 directly regulates the expression of a broad range of genes for xylem vessel formation. The Plant Journal, 66(4):579–590, 2011.

[58] Siobhan M. Brady, David A. Orlando, Ji-Young Lee, Jean Y. Wang, Jeremy Koch, José R. Dinneny, Daniel Mace, Uwe Ohler, and Philip N. Benfey. A high-resolution root spatiotemporal map reveals dominant expression patterns. Science, 318(5851):801–806, 2007.

[59] Tore Brembu, Per Winge, and Atle Magnar Bones. The small GTPase AtRAC2/ROP7 is specifically expressed during late stages of xylem differentiation in Arabidopsis. Journal of Experimental Botany, 56(419):2465–2476, 2005.

[60] Bas Jacobs, Jaap Molenaar, and Eva E. Deinum. Robust banded protoxylem pattern formation through microtubule-based directional ROP diffusion restriction. Journal of Theoretical Biology, 502:110351, 2020.

[61] Ying Zhang and Juan Dong. Cell polarity: compassing cell division and differentiation in plants. Current Opinion in Plant Biology, 45:127–135, 2018. Cell signalling and gene regulation.

[62] Yoshinobu Mineyuki. The preprophase band of microtubules: Its function as a cytokinetic apparatus in higher plants. volume 187 of International Review of Cytology, pages 1 –49. Academic Press, 1999.

[63] Pantelis Livanos and Sabine Müller. Division plane establishment and cytokinesis. Annual Review of Plant Biology, 70(1):239–267, 2019. PMID: 30795703.

[64] Yoichiro Mori, Alexandra Jilkine, and Leah Edelstein-Keshet. Wave-pinning and cell polarity from a bistable reaction-diffusion system. Biophysical Journal, 94(9):3684 – 3697, 2008.

[65] Yoichiro Mori, Alexandra Jilkine, and Leah Edelstein-Keshet. Asymptotic and bifurcation analysis of wave-pinning in a reaction-diffusion model for cell polarization. SIAM Journal on Applied Mathematics, 71(4):1401–1427, 2011.

[66] Jian-Geng Chiou, Samuel A. Ramirez, Timothy C. Elston, Thomas P. Witelski, David G. Schaeffer, and Daniel J. Lew. Principles that govern competition or co-existence in Rho-GTPase driven polarization. PLoS Computational Biology, 14(4):e1006095, 04 2018.

[67] Nicolas Verschueren and Alan Champneys. A model for cell polarization without mass conservation. SIAM Journal on Applied Dynamical Systems, 16(4):1797–1830, 2017.

[68] Corinne A Tovey and Paul T Conduit. Microtubule nucleation by γ-tubulin complexes and beyond. Essays in Biochemistry, 62(6):765–780, 2018.

[69] Akanksha Thawani, Howard A Stone, Joshua W Shaevitz, and Sabine Petry. Spatiotemporal organization of branched microtubule networks. Elife, 8:e43890, 2019.

[70] Eva Dvořák Tomaštíková, Twan Rutten, Petr Dvořák, Alisa Tugai, Klara Ptošková, Beáta Petrovská, Daniël Van Damme, Andreas Houben, Jaroslav Doležel, and Dmitri Demidov. Functional divergence of microtubule-associated tpx2 family members in arabidopsis thaliana. International journal of molecular sciences, 21(6):2183, 2020.

[71] Bandan Chakrabortty. Plant cortical microtubule dynamics and cell division plane orientation. PhD thesis, Wageningen University, Wageningen, 2017.

[72] Angela Kirik, David W. Ehrhardt, and Viktor Kirik. TONNEAU2/FASS regulates the geometry of microtubule nucleation and cortical array organization in interphase Arabidopsis cells. The Plant Cell Online, 24(3):1158–1170, 2012.

[73] J. Crank and P. Nicolson. A practical method for numerical evaluation of solutions of partial differential equations of the heat-conduction type. Mathematical Proceedings of the Cambridge Philosophical Society, 43(1):50––67, 1947.

[74] Felix Ruhnow, David Zwicker, and Stefan Diez. Tracking single particles and elongated filaments with nanometer precision. Biophysical Journal, 100(11):2820–2828, June 2011.

