## Supplementary material for "Nucleation complex behaviour is critical for cortical microtubule array homogeneity and patterning": all supplementary material

### Supplementary information for: How nucleation complexes ensure homogeneity in the plant cortical microtubule array

### 1 Supplementary text

#### 1.1 Steady state densities for non-interacting microtubules

The average microtubule density at steady state $\rho^\star$ can be calculated from the theoretically derived microtubule length distribution:

$$\rho^\star = \int_{l=0}^{l=\infty} l \cdot m(l) dl \tag{1}$$

where $m(l)$ is the density of microtubules with length $l$. Following the procedure described by [1] with the addition of the treadmilling speed, we obtain the following expression for the steady state length distributions for non-interacting growing and shrinking microtubules:

$$\begin{aligned} m(l) = m^+(l) + m^-(l) &= \frac{r_n}{v^+ - v^{tm}} e^{-\lambda l} + \frac{r_n}{v^- + v^{tm}} e^{-\lambda l} \\ &= r_n \frac{v^+ + v^-}{(v^+ - v^{tm})(v^- + v^{tm})} e^{-\lambda l}, \end{aligned} \tag{2}$$

where $r_n$ is the nucleation rate, $v^+$, $v^-$, and $v^{tm}$ are the microtubule growth, shrinkage, and treadmilling speeds, respectively, and exponent $\lambda$ is given by:

$$\lambda = \frac{r_c}{v^+ - v^{tm}} - \frac{r_r}{v^- + v^{tm}}, \tag{3}$$

where $r_c$ is the catastrophe rate and $r_r$ is the rescue rate.

Using this expression for $m(l)$, equation 1 can be solved using integration by parts:

$$\begin{aligned} \rho^\star &= \int_{l=0}^{l=\infty} l \cdot r_n \frac{v^+ + v^-}{(v^+ - v^{tm})(v^- + v^{tm})} e^{-\lambda l} dl \\ &= r_n \frac{(v^+ + v^-)(v^+ - v^{tm})(v^- + v^{tm})}{(r_c(v^- + v^{tm}) - r_r(v^+ - v^{tm}))^2}. \end{aligned} \tag{4}$$

Using this formula, we can now calculate the steady state ratio between microtubule densities in gap and band regions ($\rho^\star_{gaps}/\rho^\star_{bands}$) using the catastrophe rates for gap and band regions respectively and default values for the other parameters. This way, we obtain ratios of 0.18, 0.070, 0.038, 0.023, and 0.016 for a factor difference in catastrophe rates of 2, 3, 4, 5, and 6, respectively. These values are consistent with the ones found in our simulations with isotropic nucleation (Fig. S1B)

#### 1.2 Calculating maximal total microtubule length and initial growth speed

For comparison purposes, we want to set the average effective growth speed to be comparable to that used in previous simulations by choosing appropriate values for $v_0^+$ and $L_{max}$. The average local growth speed $\bar{v^+}$ can be obtained from equation 4 in the main text as follows:

$$\bar{v^+} = v_0^+ \frac{\bar{c_T}}{L_{max}/A}, \tag{5}$$

where $\bar{c_T}$ is the average (GTP-)tubulin concentration. This concentration can be calculated from the total tubulin length in microtubules $L_{MT}$ as follows, assuming the GDP-tubulin concentration is low compared to the GTP-tubulin concentration for models with two tubulins:

$$\bar{c_T} = \frac{L_{max} - L_{MT}}{A} = \frac{L_{max} - \sum_{j=1}^{N_{MT}} l_j}{A}, \tag{6}$$

where $N_{MT}$ is the total number of microtubules and $l_j$ is the length of the $j^{th}$ microtubule. This means the average growth speed will be given by:

$$\bar{v^+} = v_0^+ \frac{L_{max} - L_{MT}}{L_{max}}. \tag{7}$$

Therefore, the average growth speed depends on the initial growth speed and the fraction of the total amount of tubulin that is part of microtubules. Assuming that we choose an $L_{max}$ that is larger than the required microtubule length, we can calculate a value for $v_0^+$ that will end up giving us the desired growth speed:

$$v_0^+ = \bar{v^+} \frac{L_{max}}{L_{max} - L_{MT}}. \tag{8}$$

However, we still need an expression for $L_{MT}$. The total microtubule length at steady state can be calculated from the theoretically derived microtubule length distribution:

$$L_{MT} = \sum_{j=1}^{N_{MT}} l_j = \int_A \int_{l=0}^{l=\infty} l \cdot m(l) dl dA = A \int_{l=0}^{l=\infty} l \cdot m(l) dl \tag{9}$$

where $m(l)$ is the density of microtubules with length $l$, as given by equation 2 from the previous section.

Using this expression for $m(l)$, equation 9 can be solved using integration by parts:

$$\begin{aligned} L_{MT} &= A \int_{l=0}^{l=\infty} l \cdot r_n \frac{v^+ + v^-}{(v^+ - v^{tm})(v^- + v^{tm})} e^{-\lambda l} dl \\ &= r_n A \frac{(v^+ + v^-)(v^+ - v^{tm})(v^- + v^{tm})}{(r_c(v^- + v^{tm}) - r_r(v^+ - v^{tm}))^2}. \end{aligned} \tag{10}$$

With this expression we can now write down an equation for the expected growth speed at steady state (using $v^+ = \bar{v^+}$):

$$\bar{v^+} = v_0^+ \frac{L_{max} - r_n A \frac{(\bar{v}^+ + v^-)(\bar{v}^+ - v^{tm})(v^- + v^{tm})}{(r_c(v^- + v^{tm}) - r_r(\bar{v}^+ - v^{tm}))^2}}{L_{max}}. \tag{11}$$

Assuming we choose an $L_{max} > L_{MT}$, the $v_0^+$ we need is given by:

$$v_0^+ = \bar{v}^+ \frac{L_{max}}{L_{max} - r_n A \frac{(\bar{v}^+ + v^-)(\bar{v}^+ - v^{tm})(v^- + v^{tm})}{(r_c(v^- + v^{tm}) - r_r(\bar{v}^+ - v^{tm}))^2}}. \tag{12}$$

Using averages in this way is probably not entirely accurate, because most microtubules are likely to end up in band regions where the tubulin concentration is expected to be lower and, correspondingly, the growth speed will be below average. However, since differences in (GTP-)tubulin concentration are not expected to be very large, this estimate should result in reasonably comparable growth speeds.

The length distribution equations were derived for a homogeneous system, while our system uses bands and gaps with different catastrophe rates. However, under the density-dependent nucleation mechanism, most microtubules and nucleations end up in the band regions, making the band catastrophe rate a reasonable approximation. Also, because the density-dependent nucleation mechanism maintains the same global nucleation rate, while only changing the location of the nucleations, we can use the same value for $r_n A$, even though $r_n$ is locally higher in microtubule dense regions and lower in regions with few microtubules.

Using our default values for the parameters we find an $L_{MT}$ of $1.22 \cdot 10^5$ $\mu m$. The value we choose for $L_{max}$ should be larger than this value. We set it to $240000$ $\mu m$, about double the total microtubule length we expect. Using this value we can calculate that we need to set our $v_0^+$ to about $0.1$ $\mu m/s$ to end up with more or less the same growth speed.

#### 1.3 Estimation of nucleation complex parameters

There are two types of events that can occur to a nucleation complex at the membrane: dissociation and nucleation. Average times until these events occur have been measured by [2] both for complexes on microtubules and complexes not on microtubules, using labelled versions of two different nucleation complex proteins (GCP2 and GCP3). From these measurements we can make rough estimates the dissociation and nucleation rates for our simulations.

For complexes not on microtubules, the average time until an event occurs, averaged over both types of events and both GCP2 and GCP3 labels (383 observations of which 9 nucleations), is:

$$\bar{t}_{event} = 10s. \tag{13}$$

The rate $\lambda$ at which events occur is therefore:

$$\lambda = \frac{1}{\bar{t}_{event}} = 0.099s^{-1}. \tag{14}$$

The fraction $f_n$ of events that turn out to be nucleations is:

$$f_n = 0.023. \tag{15}$$

Since this number is based on only nine observations, it is very uncertain, but it should at least give us a rough estimate of the order of magnitude. From this estimate, the individual rates for nucleation and dissociation can be calculated:

$$\begin{aligned} r_{n,unbound} &= \lambda \cdot f_n = 0.0023s^{-1} \\ r_{d,unbound} &= \lambda \cdot (1 - f_n) = 0.097s^{-1}, \end{aligned} \tag{16}$$

where $r_{n,unbound}$ and $r_{d,unbound}$ are the estimated nucleation and dissociation rates for unbound complexes, respectively.

In the same way, we can calculate these parameters for microtubule-bound complexes, which have an average event time of $7.2s$ and nucleations make up a fraction of $0.24$ (based on 624 nucleations out of 2555 events). These numbers yield the following values for the bound nucleation and dissociation rates:

$$\begin{aligned} r_{n,bound} &= 0.034s^{-1} \\ r_{d,bound} &= 0.10s^{-1}, \end{aligned} \tag{17}$$

where $r_{n,bound}$ and $r_{d,bound}$ are the estimated nucleation and dissociation rates for bound complexes, respectively.

The estimated dissociation rates for bound and unbound complexes are very similar. Therefore, we used a single dissociation rate of $r_d = 0.1s^{-1}$ for all our simulations. For the nucleation rates, however, there is approximately a factor fifteen difference between estimated rates of nucleation for bound and unbound complexes. Therefore, we used a bound nucleation rate of $r_{n,bound} = 0.03s^{-1}$ and an unbound nucleation rate of $r_{n,unbound} = 0.002s^{-1}$.

Finally, we estimated a membrane association rate ($r_b$) for nucleation complexes such that the final overall nucleation rate would be close to that of $r_n = 0.001\mu m^{-2}s^{-1}$ used in the other simulations. To achieve this, we first need to estimate a total number of nucleation complexes. A nucleation complex density of 0.037 particles per $\mu m^2$ was reported by [2]. Using our total domain area, we find that on average the total number of particles $N_{NC}$ on our domain should be approximately:

$$N_{NC} = 0.037 \cdot H \cdot W = 105. \tag{18}$$

This is includes particles that have already had a nucleation but were not yet released. Those particles are not considered in our simulations, so this number represents an upper bound.

For a lower bound on this number, we can work backwards from the overall nucleation rate used in the previous simulations. If we turn this value into a rate per cell, we get:

$$r_{n,cell} = r_n \cdot A = 0.001 \cdot 2826 = 2.826 \text{ nucleations/cell/s}, \tag{19}$$

where $A$ is the area of the simulation domain. This number includes both nucleations of microtubule-bound complex and free complex. In our model context, $r_{n,cell}$ can be calculated from the total number of complexes available for nucleation, the fraction of complexes that is microtubule-bound ($f_{bound}$) and the bound and unbound nucleation rates calculated above:

$$r_{n,cell} = f_{bound} \cdot N_{NC} \cdot r_{n,bound} + (1 - f_{bound}) \cdot N_{NC} r_{n,unbound}. \tag{20}$$

We can rewrite this to estimate the number of nucleation complexes for a given nucleation rate:

$$N_{NC} = \frac{r_{n,cell}}{f_{bound} \cdot r_{n,bound} + (1 - f_{bound}) \cdot r_{n,unbound}}. \tag{21}$$

The fraction of bound complex can be estimated from the total number of events that were observed on microtubules (2555 for both GCP labels and event types out of 2938 total events) by [2], resulting in:

$$f_{bound} = 0.87. \tag{22}$$

This is not entirely accurate since we need the bound fraction at any given time and this is the fraction of observed events where the nucleation complexes were microtubule-bound. Since bound complexes have slightly faster event rates (due to their higher chance of nucleation) there will be fewer of them in our simulations at any given time than this estimate would suggest. Because the bound nucleation rate is much higher than the unbound rate, the number of complexes estimated for a given overall nucleation rate would be an underestimate. This suggests the following lower bound on the number of nucleation complexes:

$$N_{NC} = \frac{2.826}{0.87 \cdot 0.0337 + 0.13 \cdot 0.00233} = 95.4. \tag{23}$$

The number of complexes available for nucleation can be used to estimate the association rate of new nucleation complexes with the membrane. At some point the association rate must be balanced by the removal and nucleation rates to reach a steady state. Taking 100 complexes, and the overall nucleation rate from equation 19, we get:

$$r_b = N_{NC} r_d + r_{n,cell} = 100 \cdot 0.1 + 2.826 = 12.826 \text{ associations}/s = 0.0045 \text{ associations}/\mu m^2/s. \tag{24}$$

Based on this estimate, we chose an association rate of $0.0045\ \mu m^{-2} s^{-1}$.

### References


[1] Marileen Dogterom and Stanislas Leibler. Physical aspects of the growth and regulation of microtubule structures. *Physical Review Letters*, 70:1347–1350, Mar 1993.

[2] Masayoshi Nakamura, David W. Ehrhardt, and Takashi Hashimoto. Microtubule and katanin-dependent dynamics of microtubule nucleation complexes in the acentrosomal Arabidopsis cortical array. *Nature Cell Biology*, 12(11):1064–1070, November 2010.

[3] René Schneider, Kris van't Klooster, Kelsey L. Picard, Jasper van der Gucht, Taku Demura, Marcel Janson, Arun Sampathkumar, Eva E. Deinum, Tijs Ketelaar, and Staffan Persson. Long-term single-cell imaging and simulations of microtubules reveal principles behind wall patterning during proto-xylem development. *Nature Communications*, 12(1):669, 2021.

[4] Eva E Deinum, Simon H Tindemans, and Bela M Mulder. Taking directions: the role of microtubule-bound nucleation in the self-organization of the plant cortical array. *Physical Biology*, 8(5):056002, 2011.

### 2 Supplementary figures and tables

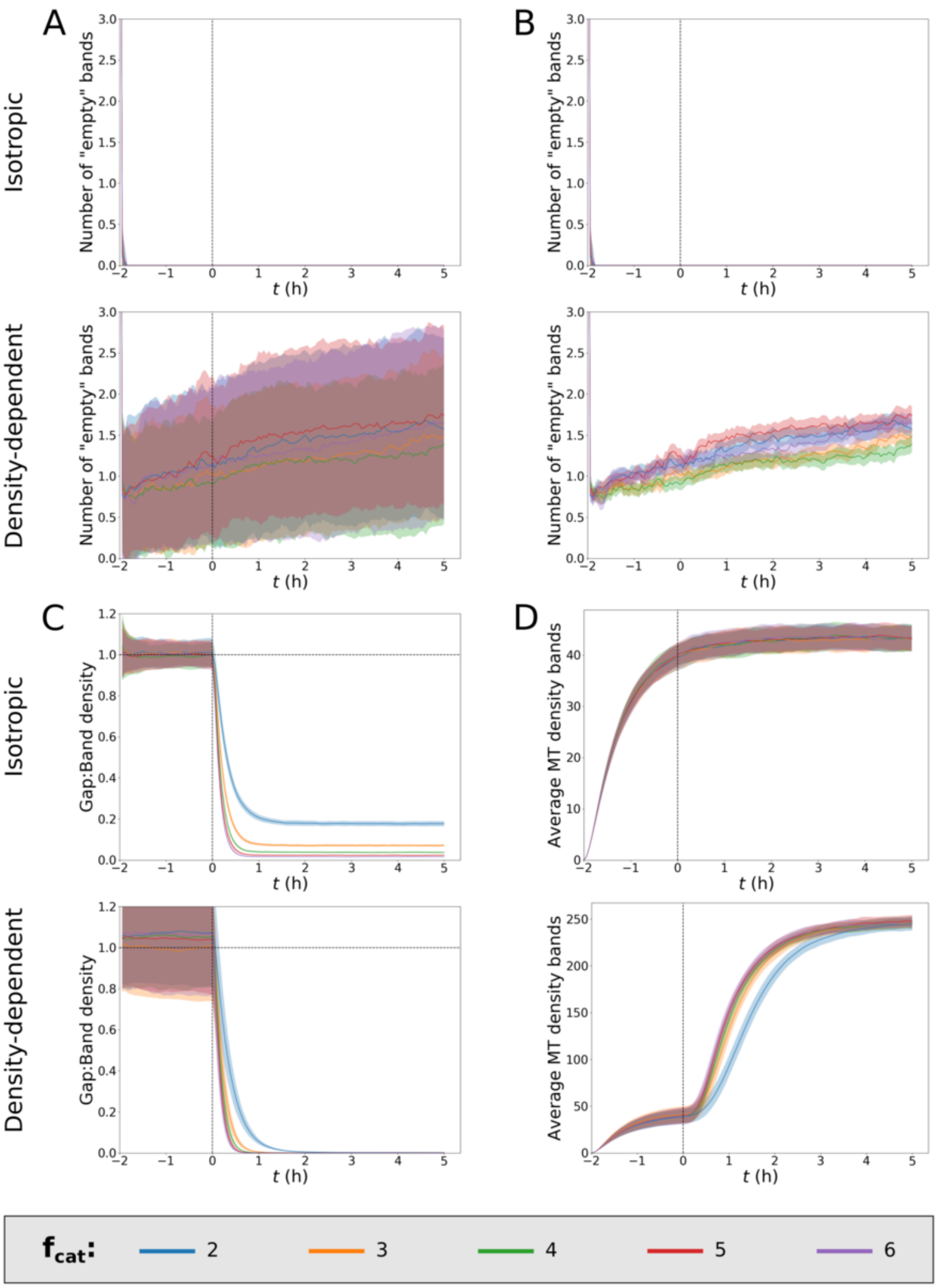


**Figure S1:** Summary quantities for protoxylem simulations with isotropic and density-dependent nucleation for various factors of difference in the catastrophe rate between bands and gaps ($f_{cat}$). (A) Number of "empty" bands (defined as bands with less than 25% of average microtubule density in bands). Note that for isotropic nucleation this quantity immediately drops to zero. (B) Same as (A) but with shaded areas representing standard error instead of standard deviation to improve readability. (C) Ratio of microtubule density between gaps and bands. (D) Average microtubule density in bands. All statistics were based on 100 simulations. Lines and shaded areas indicate mean and standard deviation (A,C,D) or standard error (B), respectively. Enhanced catastrophe rates in gap regions start at $t = 0$ h for all simulations.

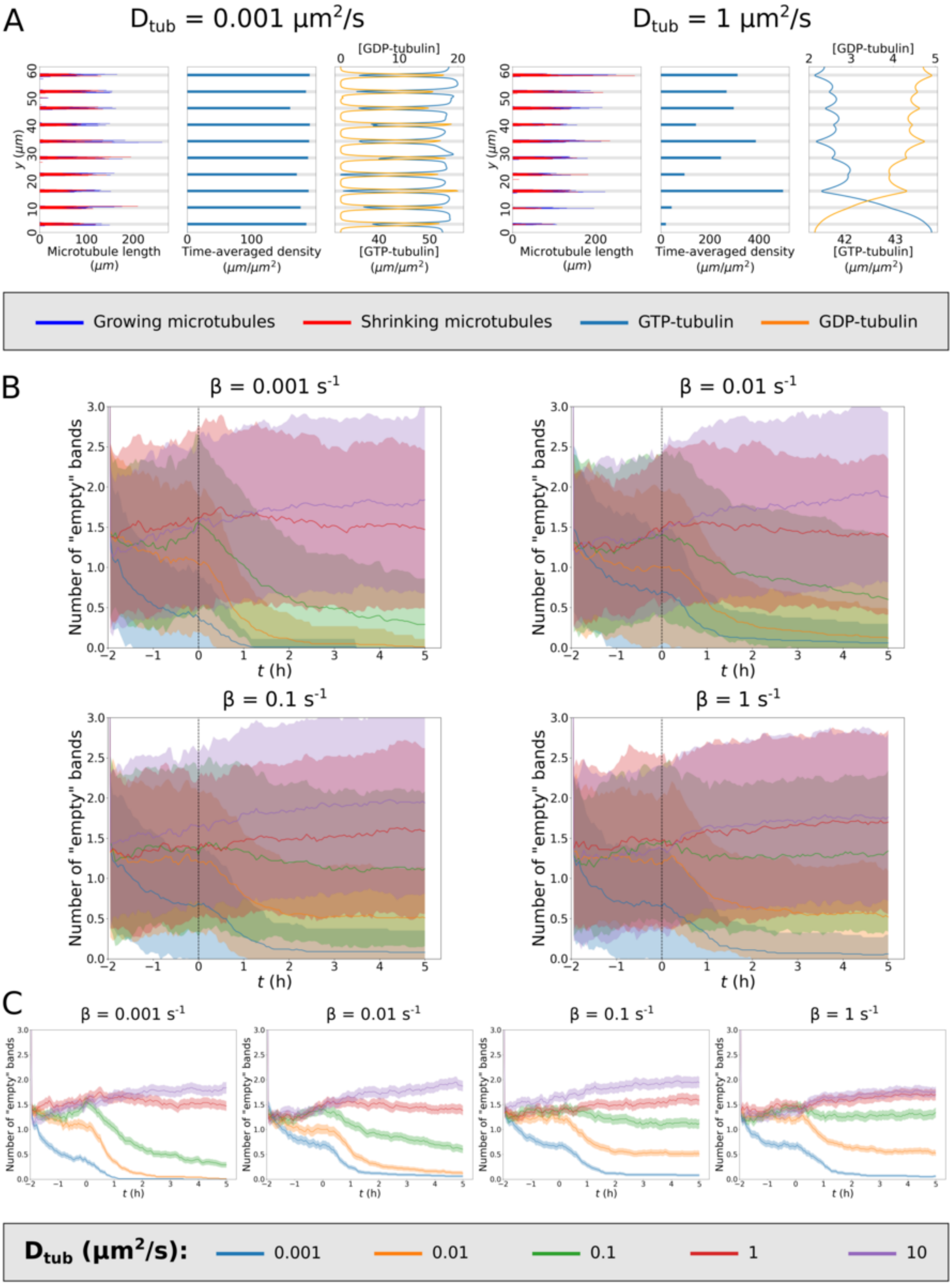


**Figure S2:** Including the extra step of GDP-tubulin to GTP-tubulin conversion facilitates formation of homogeneously populated bands, but only for biologically unrealistic parameter values. (A) Results of representative simulations with various tubulin diffusion coefficients ($D_{tub}$) for tubulin recharge rate $\beta = 0.01$. (B) Numbers of empty bands for simulations with various tubulin diffusion coefficients and recharge rates. (C) Same as (B) but with shaded areas representing standard error instead of standard deviation to improve readability. All statistics were based on 100 simulations. Lines and shaded areas indicate mean and standard deviation (B) or standard error (C), respectively.

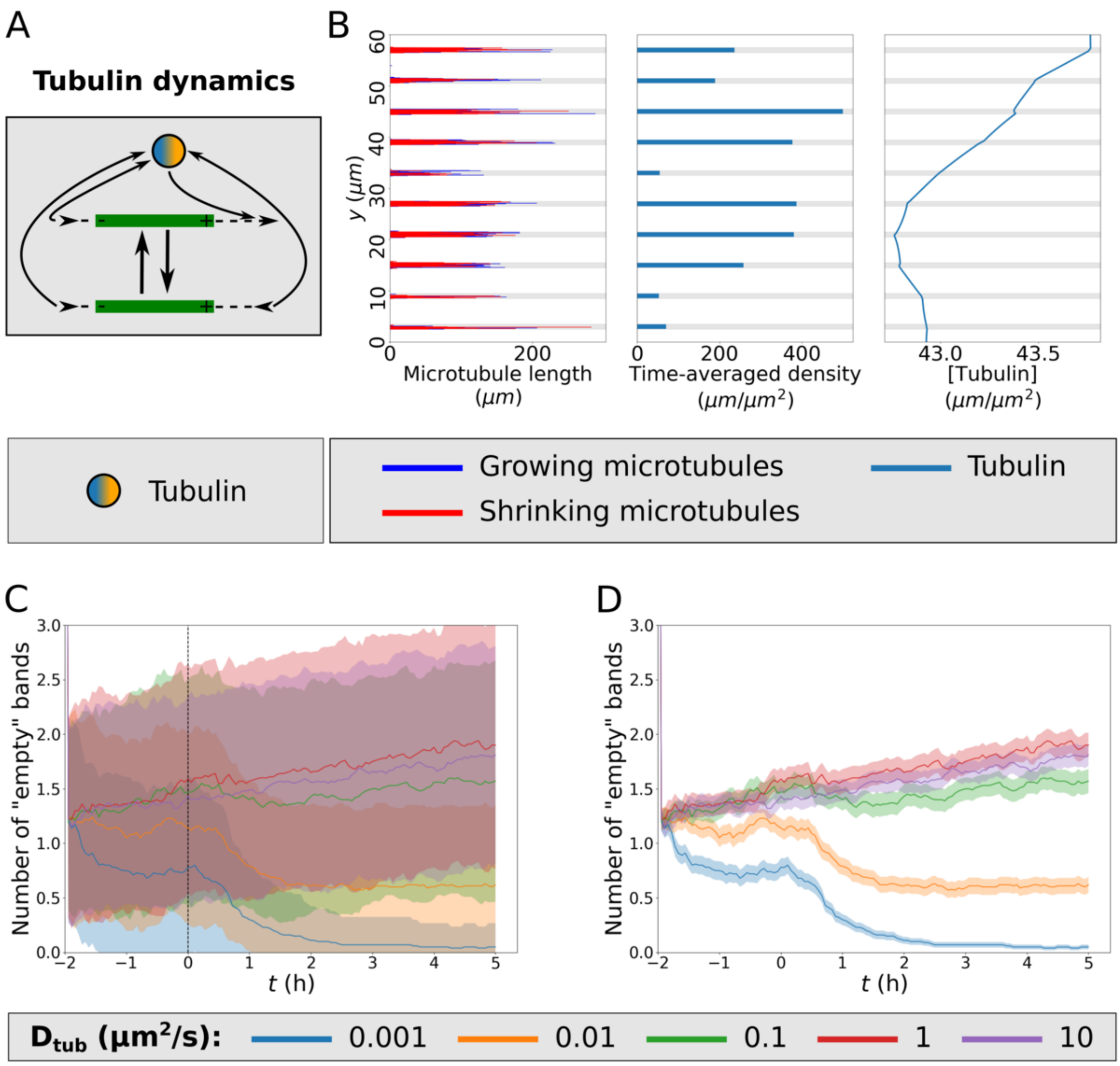


**Figure S3:** When no distinction is made between GTP- and GDP-tubulin, homogeneously populated bands are hard to achieve. (A) In our tubulin implementation, the $v^+$ of the growing microtubules depends on the local tubulin concentration, which is depleted as a result. Conversely, microtubule shrinkage increases the local tubulin concentration. (B) Results of a representative simulation with a tubulin diffusion coefficient of 1 $\mu m^2/s$. (C) Number of empty bands (bands with less than 25% of average microtubule density in bands) for simulations with various tubulin diffusion coefficients. (D) Same as (C) but with shaded areas representing standard error instead of standard deviation to improve readability. All statistics were based on 100 simulations. Lines and shaded areas indicate mean and standard deviation (C) or standard error (D), respectively.

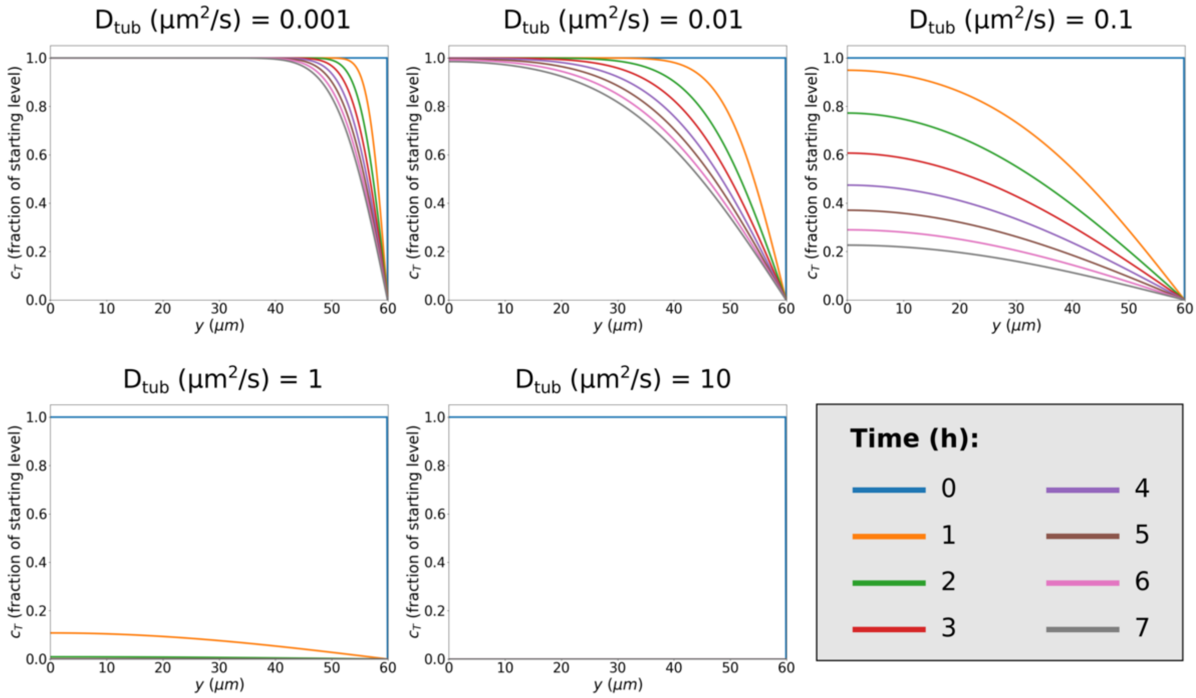


**Figure S4:** Tubulin barely diffuses from one side of the cell to the other for the lowest diffusion coefficients used in our simulations. Analytic diffusion equation solutions show the tubulin concentration ($c_T$) at one hour intervals for various tubulin diffusion coefficients ($D_{tub}$) in absence of microtubules. The left boundary had zero flux boundary conditions and the right boundary was fixed at zero to provide a maximally effective tubulin sink.

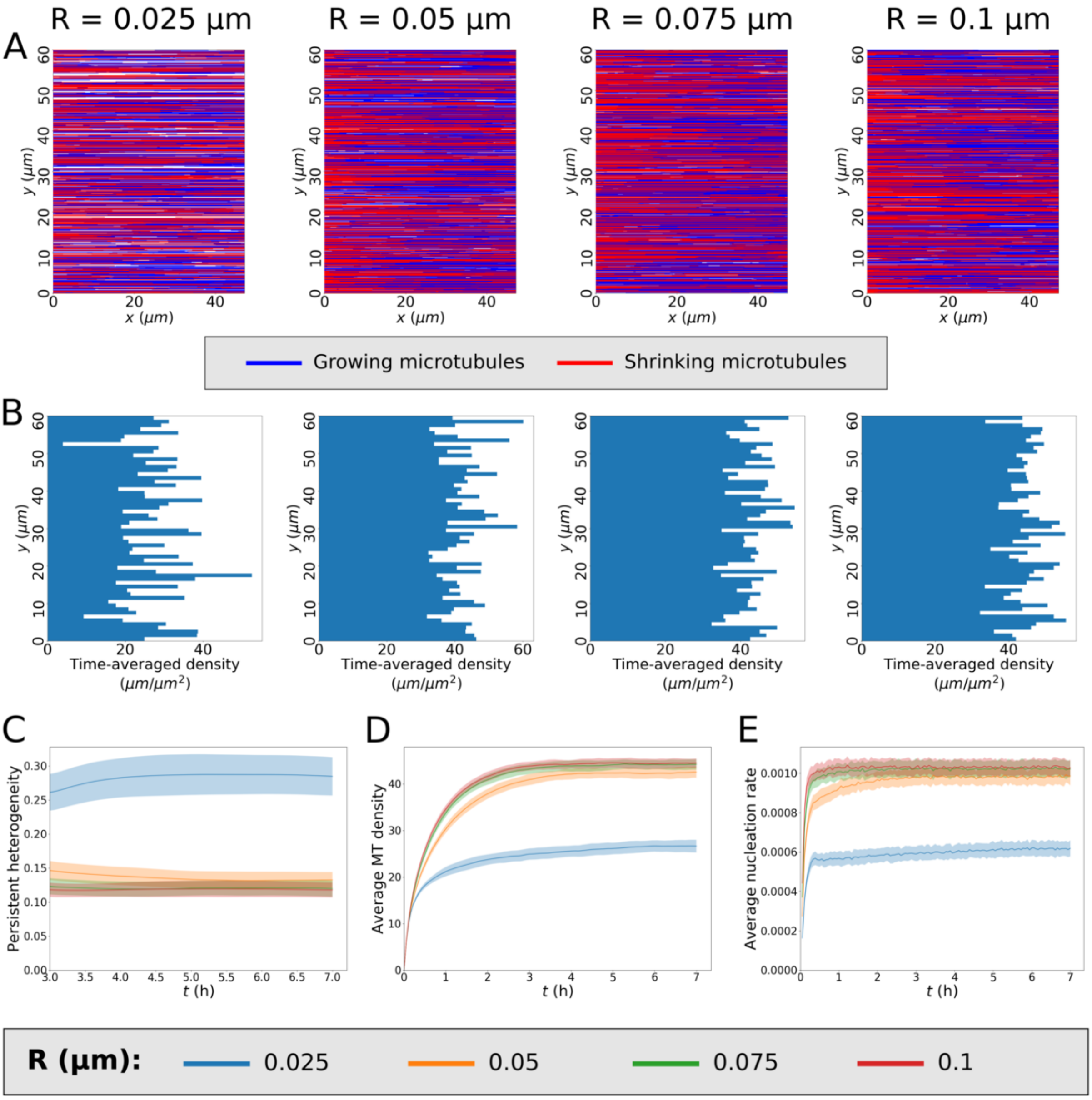


**Figure S5:** With microtubule-dependent nucleation complex insertion, homogeneous arrays can be established as long as the radius $R$ at which microtubules attract complexes is sufficiently large. (A) Results of representative simulations with different values of $R$. (B) Time-averaged microtubule densities over the last three hours of the corresponding simulations from (A). (C) Measure of persistent homogeneity. (D) Average microtubule density of the entire array over time. (E) Average global nucleation rate (nucleations$/\mu m^2/s$). All statistics were based on 100 simulations with $D_{NC} = 0.013\ \mu m^2/s$ and $r_{b,min} = r_b/40\ \mu m^{-2}s^{-1}$. Lines and shaded areas indicate mean and standard deviation, respectively.

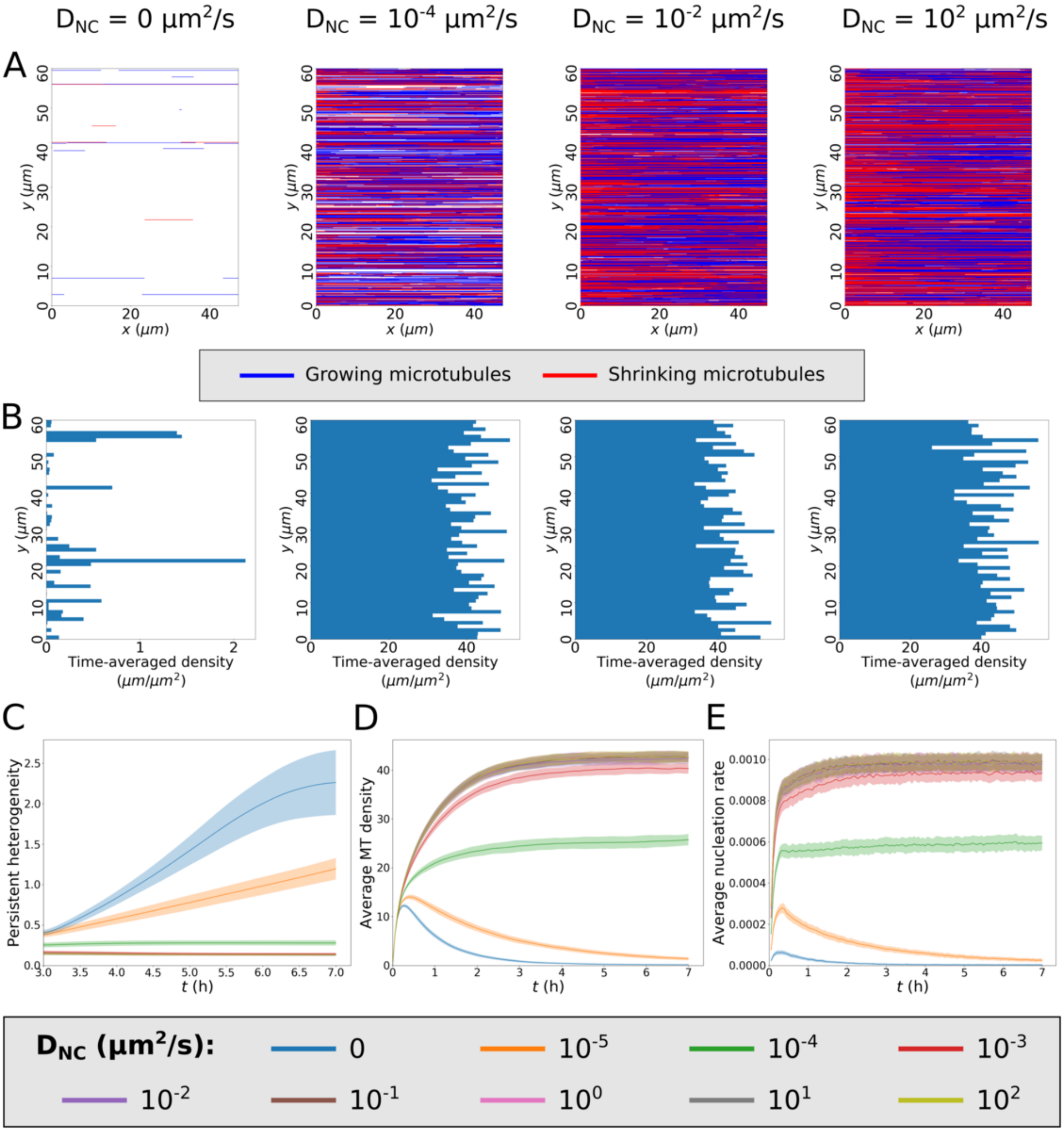


**Figure S6:** The nucleation complex implementation with microtubule-dependent insertion and a reduced nucleation rate for unbound complex yields homogeneous arrays as long as diffusion coefficients are high enough for complexes to find and bind microtubules. (A) Results of representative simulations with different nucleation complex diffusion coefficients ($D_{NC}$). (B) Time-averaged microtubule densities over the last three hours of the corresponding simulations from (A). (C) Measure of persistent homogeneity. (D) Average microtubule density of the entire array over time. (E) Average global nucleation rate (nucleations/$\mu m^2/s$). All statistics were based on 100 simulations with $R = 0.05\ \mu m$ and $r_{b,min} = r_b/40\ \mu m^{-2}s^{-1}$. Lines and shaded areas indicate mean and standard deviation, respectively.

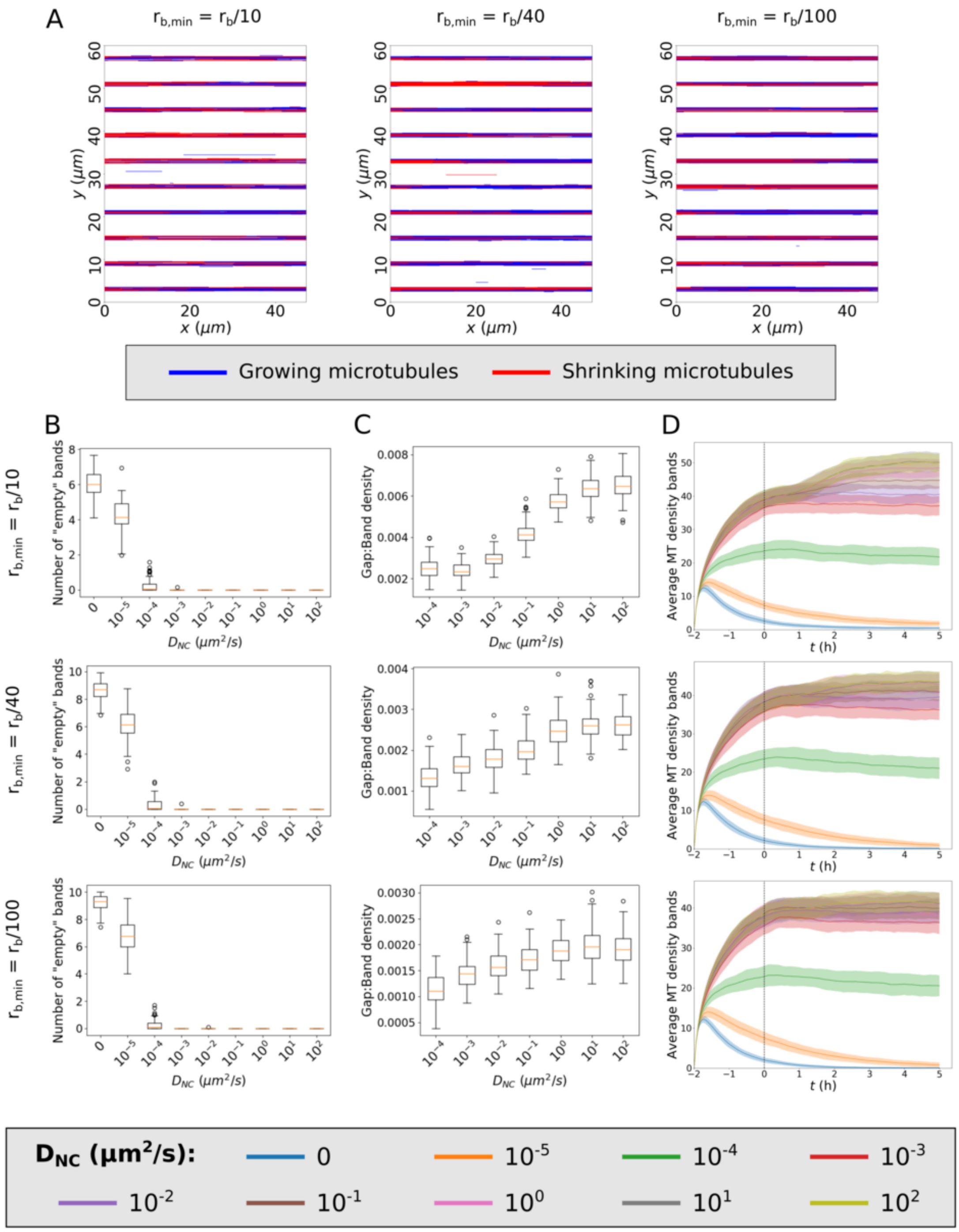


**Figure S7:** Microtubule-dependent nucleation complex insertions greatly improves band formation without resulting in empty bands for reasonable parameter values. (A) Overview of microtubule coverage of the cell membrane for representative simulations with a factor 3 difference in catastrophe rates between bands and gaps, using various values of $r_{b,min}$, and a $D_{NC}$ of $0.013\ \mu m^2/s$. (B) Average numbers of empty bands (bands with less than 25% of average microtubule density in bands) during the last two hours of the simulation. (C) Average ratios between microtubule density in gaps and bands during the last two hours of the simulations. The lowest two diffusion coefficients were disregarded here, because a lack of array density resulted in zero divisions. (D) Microtubule density in bands over time. All statistics were based on 100 simulations for various values of $D_{NC}$ and $r_{b,min}$. Lines and shaded areas indicate mean and standard deviation, respectively. Enhanced catastrophe rates in gap regions start at $t = 0$ h for all simulations.

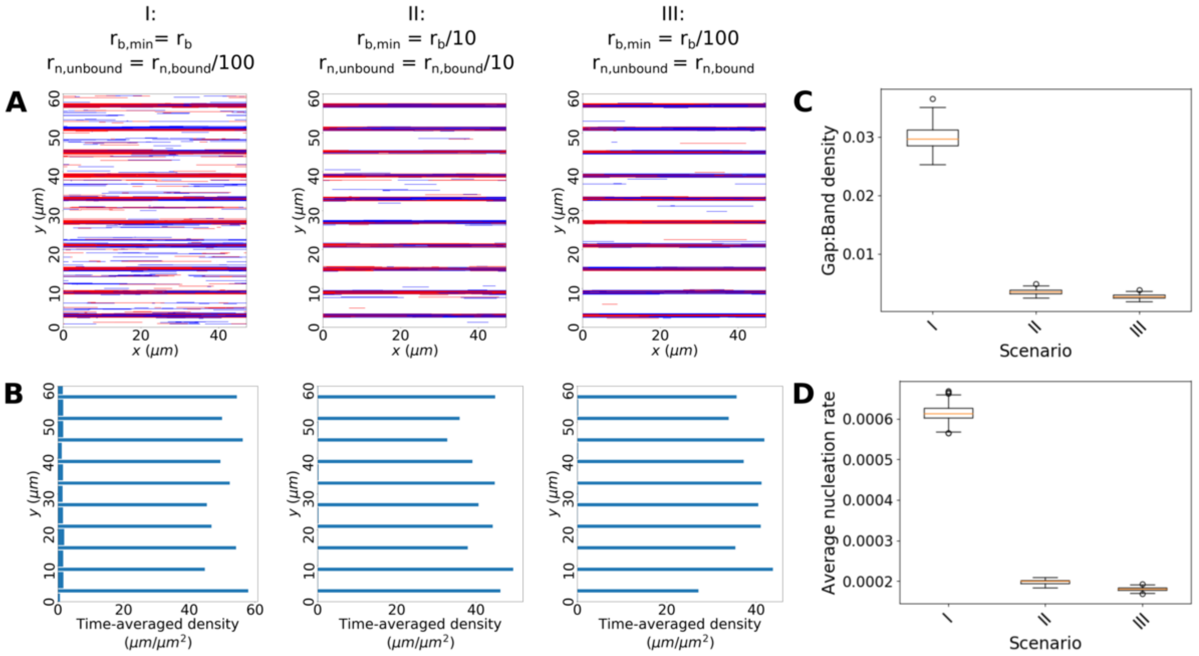


**Figure S8:** Microtubule-dependent nucleation complex insertion is more effective at improving protoxylem band formation than increased nucleation for microtubule-bound complexes. (A) Representative simulations using a factor 3 difference in catastrophe rates between bands and gaps and $D_{NC} = 0.013 \mu m^2/s$ for various nucleation and insertion scenarios: I: No reduction in insertion rate ($r_b$) in absence of microtubules and a hundred fold reduction in nucleation rate ($r_n$) for unbound complexes. II: A ten fold reduction in the insertion rate without nearby microtubules and a ten fold reduction in nucleation rate for unbound complexes. III: A hundred fold reduction in the insertion rate without nearby microtubules and no difference in nucleation rate for unbound complexes. (B) Time-averaged microtubule densities over the last three hours of the corresponding simulations from (A). (C) Average ratios between microtubule density in gaps and bands during the last two hours of the simulations. (D) Average global nucleation rate during the last two hours of the simulations. All statistics were based on 100 simulations for the three scenarios described in (A). Enhanced catastrophe rates in gap regions start at $t = 0$ h after two hours without difference in catastrophe rates and end at $t = 5$ h for all simulations.

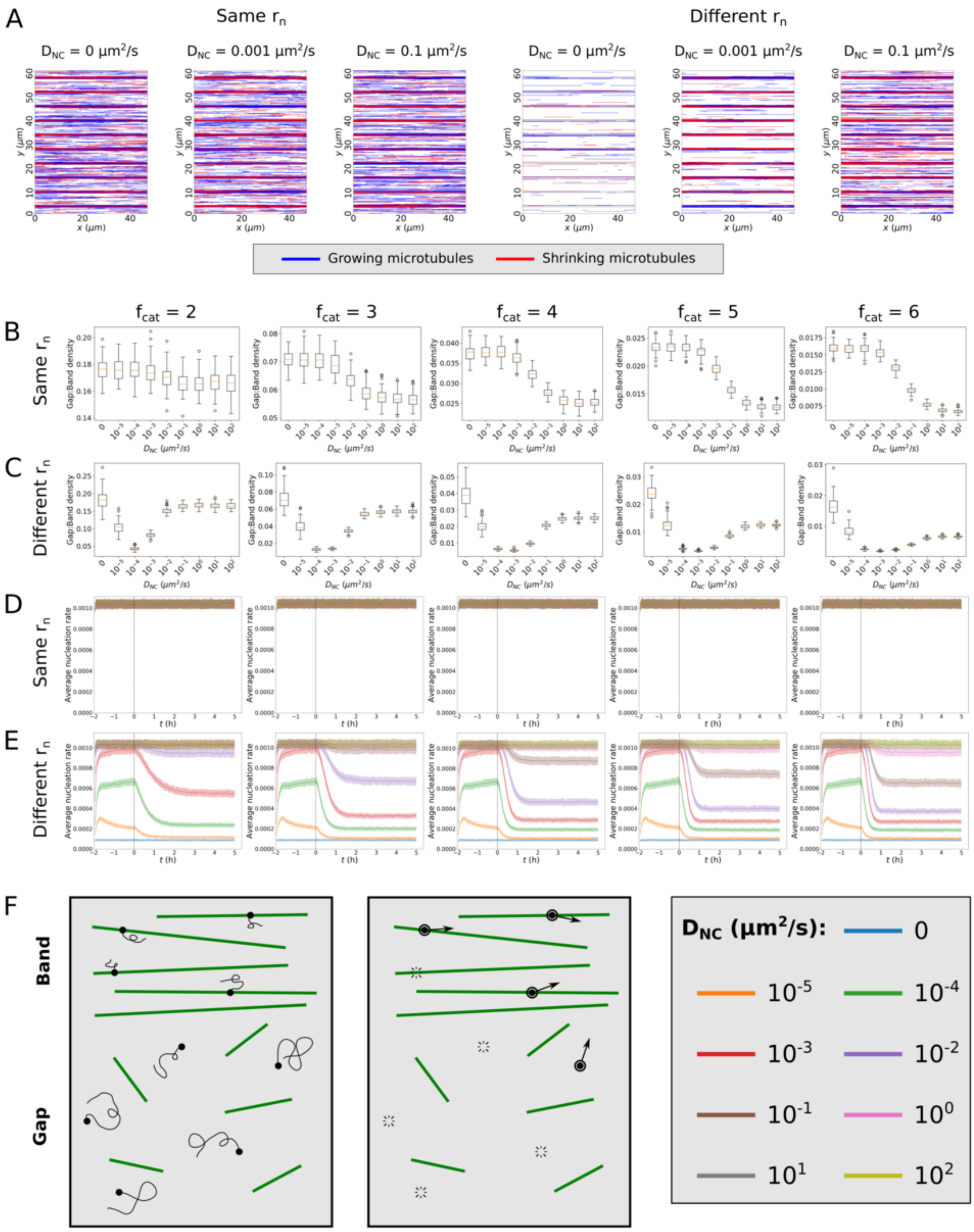


**Figure S9:** In absence of microtubule-dependent nucleation complex insertion, a reduced nucleation rate for unbound complexes results in an optimum in the ratio between gap and band density for intermediate values of the nucleation complex diffusion coefficient ($D_{NC}$). (A) Representative simulations with and without a different nucleation rate for microtubule-bound and unbound nucleation complexes using a factor 3 difference in catastrophe rates between bands and gaps and various values of $D_{NC}$. (B,C) Average ratios between microtubule density in gaps and bands during the last two hours of the simulations with identical (B) and reduced (C) $r_n$ in gaps. Note that –coming from high $D_{NC}$–, the window with lowest gap:band density ratio (i.e., optimal separation) in $C$ starts from $D_{NC}$ values for which the total microtubule density decreases at the start of the initiation phase. (D,E) Average global nucleation rates over time. All statistics were based on 100 simulations with (C,E) and without (B,D) a reduced nucleation rate for unbound complexes, for various factors of difference in band and gap catastrophe rates ($f_{cat}$) and various nucleation complex diffusion coefficients ($D_{NC}$). Lines and shaded areas indicate mean and standard deviation, respectively. Enhanced catastrophe rates in gap regions start at $t = 0$ h for all simulations. (F) Proposed mechanism of separation enhancement. Nucleation complexes in the bands are likely to encounter a microtubule in their lifetime and will therefore benefit from the higher bound nucleation rate, while complexes in the gaps are more likely to disappear before they encounter a microtubule and therefore largely follow the lower unbound nucleation rate.

**Table S1:** Nucleation complexes insert more readily near existing microtubules. Comparison of nucleation complex insertions in oryzalin-treated cells and controls. The literature estimate for the insertion rate that we used in the simulations to keep the overall nucleation rate comparable was 0.0045 $\mu m^{-2} s^{-1}$.

| Treatment:<br>Area: | Mock<br>Total | Oryzalin<br>Total | Oryzalin<br>Non-microtubule | Oryzalin<br>Microtubule | Unit |
|---|---|---|---|---|---|
| Number of nucleation complexes inserted | 338 | 170 | 55 | 115 | - |
| Membrane area | 873 | 5445 | 5372 | 73 | $\mu m^2$ |
| Measurement time | 120 | 120 | 120 | 120 | $s$ |
| Estimated insertion rate | 0.0037 | 0.00026 | 0.000085 | 0.013 | $\mu m^{-2} s^{-1}$ |
| Factor lower than literature estimate | 1.4 | 17.3 | 52.7 | 0.35 | - |

**Table S2:** Default parameter values used in the simulations.

| Parameter | Value | Unit | Description | Source |
|---|---|---|---|---|
| Simulation domain | | | | |
| $H$ | 60 | $\mu m$ | Domain height | [3] |
| $W$ | $7.5 \cdot 2\pi$ | $\mu m$ | Domain circumference | [3] |
| Standard parameters | | | | |
| $v^+$ | 0.05 | $\mu m/s$ | Growth speed | [3] |
| $v^-$ | 0.08 | $\mu m/s$ | Shrinkage speed | [3] |
| $v^{tm}$ | 0.01 | $\mu m/s$ | Treadmilling speed | [3] |
| $r_c$ | 0.0016 | $s^{-1}$ | Catastrophe rate (bands) | [3] |
| $r_r$ | 0.001 | $s^{-1}$ | Rescue rate | [3] |
| $r_n$ | 0.001 | $\mu m^{-2} s^{-1}$ | Nucleation rate | [3] |
| $f_{cat}$ | 3 | | Factor increased catastrophe rate in gaps | |
| Density-dependent nucleation | | | | |
| $\rho_{\frac{1}{2}}$ | 0.1 | $\mu m/\mu m^2$ | Microtubule density at which half of all nucleations are microtubule-bound | [4] |
| Tubulin limited growth | | | | |
| $D_{tub}$ | Variable | $\mu m^2/s$ | Tubulin diffusion coefficient | |
| $\beta$ | 0.01 | $s^{-1}$ | Tubulin recharge rate | |
| $v_0^+$ | 0.1 | $\mu m/s$ | Initial growth speed | Appendix 1.2 |
| $L_{max}$ | $2.4 \cdot 10^4$ | $\mu m$ | Maximum total microtubule length | Appendix 1.2 |
| Nucleation complexes | | | | |
| $r_b$ | 0.0045 | $\mu m^{-2} s^{-1}$ | Insertion rate | Appendix 1.3 |
| $r_d$ | 0.1 | $s^{-1}$ | Dissociation rate | Appendix 1.3 |
| $D_{NC}$ | 0.013 | $\mu m^2/s$ | Nucleation complex diffusion coefficient | This paper |
| $r_{n,bound}$ | 0.03 | $s^{-1}$ | Bound nucleation rate | Appendix 1.3 |
| $r_{n,unbound}$ | 0.002 | $s^{-1}$ | Unbound nucleation rate | Appendix 1.3 |
| Microtubule-dependent nucleation complex insertion | | | | |
| $r_{b,min}$ | $r_b$/40 | $\mu m^{-2} s^{-1}$ | Insertion rate in absence of microtubules | |
| $R$ | 0.05 | $\mu m$ | Radius in which nucleation complex insertion is enhanced by microtubules | |
